# A non-spiking neuron model with dynamic leak to avoid instability in recurrent networks

**DOI:** 10.1101/2020.06.16.155614

**Authors:** Udaya B. Rongala, Jonas M.D. Enander, Matthias Kohler, Gerald E. Loeb, Henrik Jörntell

## Abstract

Recurrent circuitry components are distributed widely within the brain, including both excitatory and inhibitory synaptic connections. Recurrent neuronal networks have potential stability problems, perhaps a predisposition to epilepsy. More generally, instability risks making internal representations of information unreliable. To assess the inherent stability properties of such recurrent networks, we tested a linear summation, non-spiking neuron model with and without a ‘dynamic leak’, corresponding to the low-pass filtering of synaptic input current by the RC circuit of the biological membrane. We first show that the output of this neuron model, in either of its two forms, follows its input at a higher fidelity than a wide range of spiking neuron models across a range of input frequencies. Then we constructed fully connected recurrent networks with equal numbers of excitatory and inhibitory neurons and randomly distributed weights across all synapses. When the networks were driven by pseudorandom sensory inputs with varying frequency, the recurrent network activity tended to induce high frequency self-amplifying components, sometimes evident as distinct transients, which were not present in the input data. The addition of a dynamic leak based on known membrane properties consistently removed such spurious high frequency noise across all networks. Furthermore, we found that the neuron model with dynamic leak imparts a network stability that seamlessly scales with the size of the network, conduction delays, the input density of the sensory signal and a wide range of synaptic weight distributions. Our findings suggest that neuronal dynamic leak serves the beneficial function of protecting recurrent neuronal circuitry from the self-induction of spurious high frequency signals, thereby permitting the brain to utilize this architectural circuitry component regardless of network size or recurrency.

**Author Summary:** It is known that neurons of the brain are extensively interconnected, which can result in many recurrent loops within its neuronal network. Such loops are prone to instability. Here we wanted to explore the potential noise and instability that could result in recurrently connected neuronal networks across a range of conditions. To facilitate such simulations, we developed a non-spiking neuron model that captures the main characteristics of conductance-based neuron models of Hodgkin-Huxley type, but is more computationally efficient. We found that a so-called dynamic leak, which is a natural consequence of the way the membrane of the neuron is constructed and how the neuron integrates synaptic inputs, provided protection against spurious, high frequency noise that tended to arise in our recurrent networks of varying size. We propose that this linear summation model provides a stable and useful tool for exploring the computational behavior of recurrent neural networks.

## Introduction

Recurrent excitatory loops are a common feature in the central nervous system, such as in neocortical circuits [1–4], thalamocortical loops [5,6], cerebrocerebellar and spinocerebellar loops [7,8]. Inhibitory interneurons have been described to provide lateral inhibition [9–12] and feed-forward inhibition [13,14], but they also make synapses on other inhibitory neurons, thereby potentially forming recurrent disinhibitory loops as well [15–17]. Furthermore, such excitatory and inhibitory connectivity has been reported to be balanced [18–20]. Functionally, recurrent connections enable a network to use preceding states to impact the processing of the present state. Such state memory can, for example, improve learning performance [21]. However, due to the many potential positive feedback loops in larger networks with extensive recurrent connections, imbalances in excitatory (E) and inhibitory (I) synaptic activity could lead to activity saturation [22,23], such as observed in epilepsy [24,25], or, in milder cases, a noise-like perturbation of the information content of internal signals, which would be disadvantageous for learning.

We explored potential noise and stability issues that could arise in recurrent neuronal networks. In order to focus on the network architecture aspect of this problem, we used a non-spiking neuron model designed to be simple and computationally efficient, while embodying fundamental properties of Hodgkin-Huxley conductance-based models. The relevance of a non-spiking neuron model stems from the stochasticity inherent in neuronal spike generation [26–28], which renders the spiking output of the individual neuron to some degree unreliable in terms of information content. To compensate for such unreliability, the brain could encode each representation across a population of neurons (below referred to as an ensemble of redundant neurons), as has been observed in the brain *in vivo* [29]. The input-output relationships across a range of neuron types in the central nervous system *in vivo* indicate that overall, each neuron’s spike output is a probability density function (PDF) of the underlying membrane potential of the neuron [27]. That PDF thereby approximates the membrane potential and could be considered to correspond to the spike output of an ensemble of neurons with similar inputs. Thus, simulating a non-spiking neuron and providing the PDF of the neuron as its output avoids the extreme resource demands of both simulating the highly complex spike generation stochasticity [28] and compensating for that stochasticity by simulating large populations of redundant neurons. Synaptic input creates modulation of the neuronal membrane potential, hence its PDF, by temporarily activating conductances that are added to the static leak conductances. The synaptic conductances and currents can modulate very rapidly but the membrane capacitance together with the static leak channels forms an RC circuit that constitutes a low-pass filter (herein, dynamic leak) for the resultant membrane potential. We hypothesized that this dynamic leak would improve network stability without compromising information transfer.

To test this hypothesis, we constructed a highly recurrent, two-layer neuronal network, with five excitatory and five inhibitory neurons in the first layer and four excitatory and four inhibitory neurons in the second layer. All neurons in both layers were reciprocally connected with randomized gains. All first layer neurons were provided with six randomized and broadly distributed input signals. A striking finding was that for all tested network configurations, synaptic weight distributions, various conduction delays and input density of sensory inputs, recurrent networks tended to generate high frequency components that were not present in the sensory input data. In all cases these transients were eliminated by incorporating a dynamic leak in the neuron models without compromising the representation of the input signals.

We note that the fully reciprocal connectivity employed in the networks described herein encompasses the wide range of connectivity that has been identified experimentally in cortical and other central neural structures (see above). The strictly layered connectivity of many popular neural network models for deep learning reflects only a small subset of the known complexity of biological networks. Attempts to add limited recurrency into such models have encountered stability problems [22,23], for which dynamic leak appears to offer substantial mitigation.

## Methods

### Neuron model

#### Linear summation neuron model (LSM)

The neuron model implemented for this study was a non-spiking, linear input summation model with an additional dynamic leak component. For the version without dynamic leak, the activity 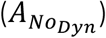 was given by the total weighted input activity (*w* ∗*a*) (where *a* is the activity of each individual input and *w* is the weight of that input) across all individual synapses (*i*) (Eq. 1). Electrotonic compactness in the neuron is assumed, so that all synapses have equal impact on the activity of the neuron. This simplified model of synaptic input activity integration can be shown to be closely related to a Hodgkin-Huxley (H-H) model (see Appendix 1), for example resulting in the preservation of two key dependencies of EPSPs and IPSPs on membrane biophysics: *i*) input response magnitude depends on the difference between the membrane potential and the reversal potentials for the relevant corresponding ‘ion channels’ (i.e. depending on if the input is provided through an excitatory or an inhibitory synapse); *ii*) input response magnitude depends on relative shunting of synaptic currents by conductances resulting from the background of synaptic input activity (Eq. 1). The responsive properties of the LSM and the H-H neuron model are shown to be highly similar in S1 Fig.

The LSM implemented a degree of activity normalization (denominator of Eq. 1) by introducing a static leak, which was calculated as the product of a constant (*k*_*static*_) multiplied by the number of synapses on the neuron, plus a term reflecting the total number of open channels, which is activity dependent.

To mimic the effect of the RC circuit created by the ion channels and the capacitance of the membrane, we added a dynamic leak function to the neuron. To test the impact of the dynamic leak on network dynamics, we compared the networks composed of neurons with the dynamic leak with the same network when the neuron model did not include this dynamic leak. The neuron activity for the neuron model variant with dynamic leak (*A*_*Dyn*_) is given by the linear summation model with an additional leak time constant (*τ*_*Dyn*_). Larger neurons with more synapses tend to have longer time constants [30], so we tried various ways of scaling *τ*_*Dyn*_ with number of synapses *i*. Thereby, the dynamic leak integrates the function of the capacitance in the RC circuit of the biological neuron into the LSM (Eq. 2). The neuron activity of this model is given by the following equations,

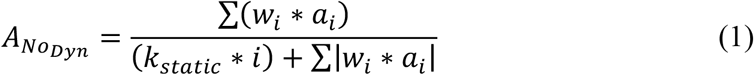

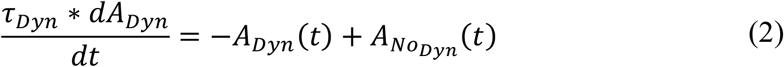

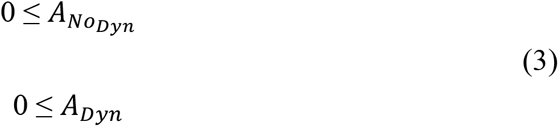

Figure 1 illustrates the output activity of individual LSM neurons (Eq. 1-3), which were isolated in the sense that they were not connected to any neuronal network other than the provided inputs, for different input combinations of emulated excitatory and inhibitory synaptic inputs (Fig 1A, B). The input spike trains were convoluted using a kernel function in order to emulate post-synaptic-potential inputs (detailed below, Eq. 6), that were fed to the LSM neuron (Fig 1C-E). The LSM activity without dynamic leak (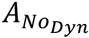, Fig 1F) shows the activity normalization resulting from the static leak constant (*k*_*static*_ = 1, for this illustration), along with the effect of the neuron output activity threshold at zero (Eq. 3). The activity of the LSM neuron would also be expected to fall back towards this zero level of activity without any external or internal input. This level hence corresponds to a threshold for spike initiation among a population of similarly connected neurons that are typically represented by the one modeled neuron. The output activity for the LSM neuron with dynamic leak (*A*_*Dyn*_, Fig 1G) exhibits a low pass filtering effect on the output activity, which is reflective of the effect of the RC component integrated in the LSM neuron model with the dynamic leak.

**Fig 1.**
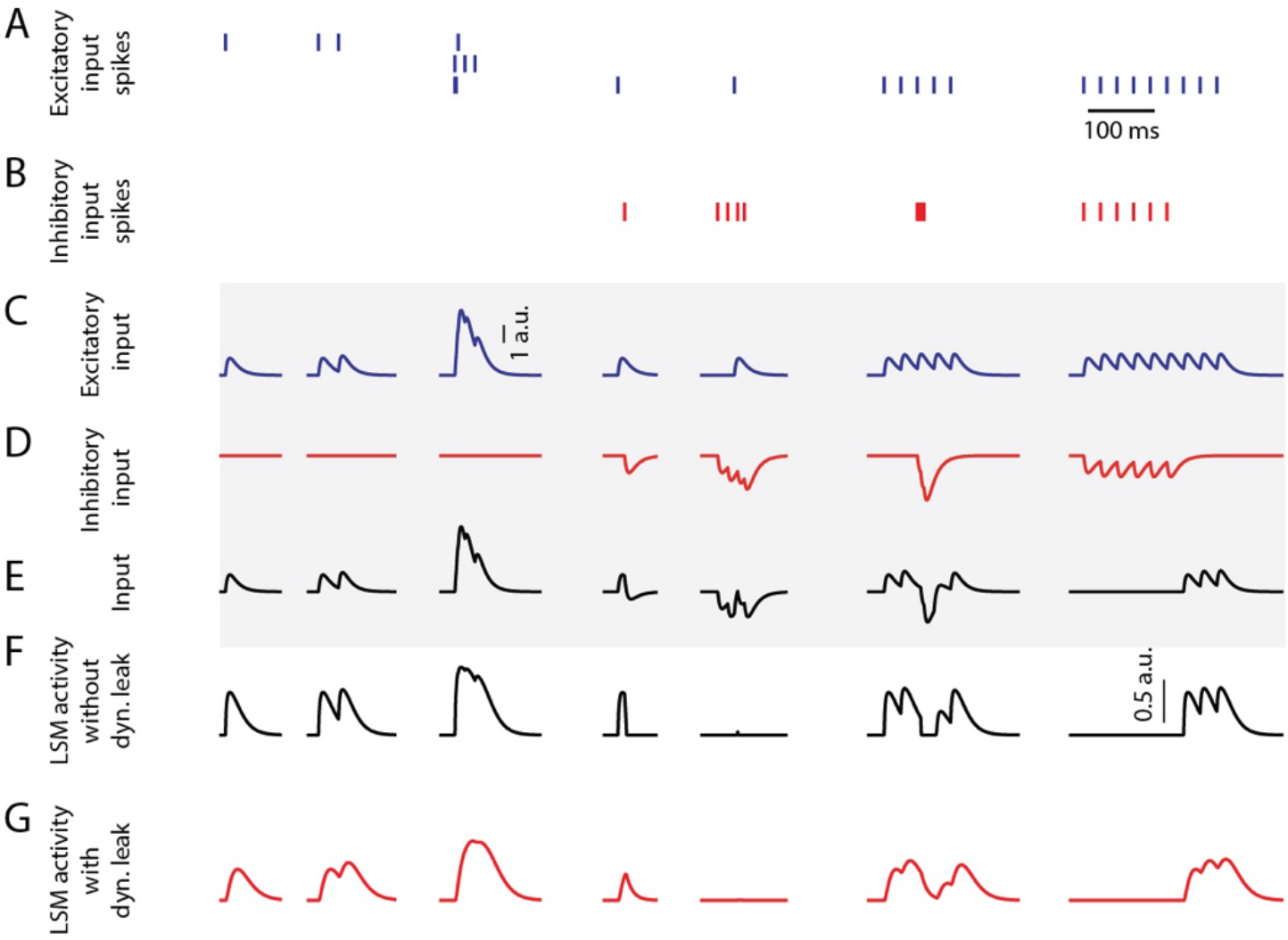
LSM responses to emulated synaptic inputs. (**A**) The activation times of three different excitatory synaptic inputs are indicated as spike trains. (**B**) The activation times of one inhibitory synaptic input. (**C-D**) The excitatory and inhibitory sensory input spike trains were convoluted using a kernel function (see Methods) to create input that resembles post-synaptic potentials. Note that the input to the LSM neuron can exceed 1 a.u., while the output of the LSM neuron cannot. Calibration applies to C –E (traces in the shaded region). (**E**) The input from summation of the excitatory and inhibitory inputs. (**F**) LSM (without dynamic leak, *k*_*static*_ = 1) output activity for the given PSP inputs. Calibration applies to F-G. (**G**) LSM (with dynamic leak, *k*_*static*_ = 1, τ_*dyn*_ = 1/100 *s*) output activity for the given PSP inputs.

Figure 2 illustrates the impact of various static and dynamic leaks. As indicated in Fig 2A, the static leak constant acts as a normalization factor for the total neuron activity, without diminishing the underlying dynamics of that activity (S2 Fig). At very low values of the static leak constant, the mean activity reached sufficiently high levels for the reversal potential to start having a significant dampening effect on the activity dynamics (see uppermost trace in Fig 2A), substantially reducing the coefficient of variation (CV in Fig 2B). Fig 2C, D illustrates the additional impact of various values of the dynamic leak constant. Fig 2C and S3 Fig demonstrate the filtering effect of the dynamic leak constant on the total neuron activity. A high value of this dynamic leak constant substantially smoothens the activity dynamics, which was reflected in the resulting low CV value (Fig 2D). The dynamic leak constant (*τ*_*Dyn*_) was set to 1/100 for the rest of this study, unless otherwise specified.

**Fig 2.**
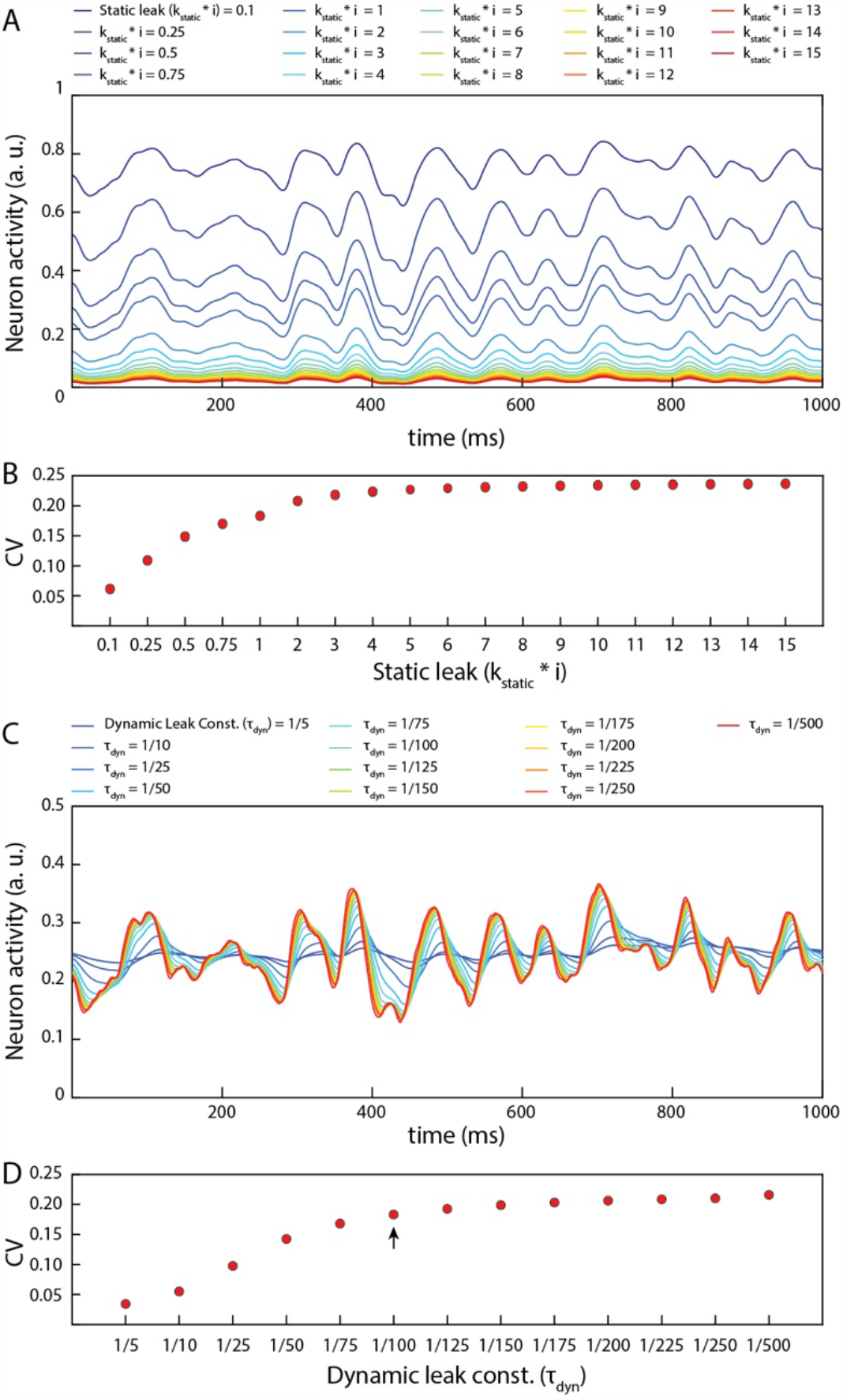
Impact of static leak (K_static_) and dynamic leak constant (τ_dyn_) in LSM. (**A**) Impact of the value of *k*_*static*_ in the LSM (for τ_*dyn*_ = 1/100) for a given pseudo-random sensory input at 50 Hz for each of six sensors (see Fig 3). (**B**) The perseverance of dynamics in the neuron activity (*A*) for varying *k*_*static*_ value as assessed by the coefficient of variation (*C*V = σ (A)/Á). A higher value of CV indicates a higher activity variance relative to the mean activity. (**C**) Impact of the value of τ_*Dyn*_ in the LSM (for *k*_*static*_ = 1) for the same pseudo-random sensory input as in **A**. (**D**) The CV as a function of the value of the dynamic leak (τ_JKL_) in the LSM. The arrow indicates the value of τ_JKL_ used in rest of this paper unless otherwise specified.

In Appendix 1, we show that the LSM neuron model can be derived from a simplified H-H type conductance-based neuron model. In the H-H model, the leak is proportional to the membrane voltage and the synaptic currents are scaled depending on the membrane voltage, so that the voltage is limited to a fixed range. The differential equation describing this model suffers from numerical instability, therefore we solve it with the implicit Euler method. The model is simple enough so that an analytical solution can be obtained. Key H-H model features that are captured by the LSM neuron: *i*) the response to a given input scales with the difference between the current activity level (membrane potential *V*) and the reversal potentials of the excitatory/inhibitory inputs (which have been normalized to +1 and −1, respectively); and *ii*) the impact of a given input is scaled by the degree of the shunting caused by the total synaptic activity the neuron receives at that time.

#### Izhikevich neuron model (IZ)

For the Izhikevich neuron model [31], the membrane potential (*IZ*_*v*_) and the adaptation variable (*IZ*_*u*_) were updated via the following nonlinear differential equations discretized using Euler’s method.

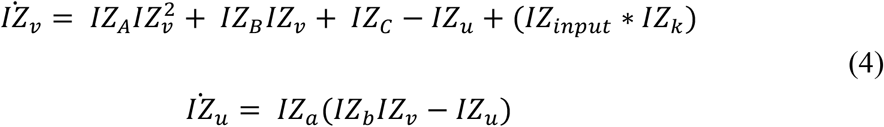

When the membrane potential reached the spike depolarization threshold of 30 *mV*, one spike was produced followed by a reset:

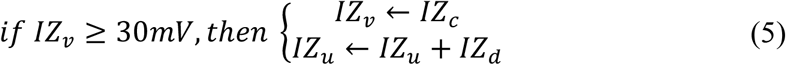

The *IZ*_*A*_, *IZ*_*B*_ & *IZ*_*C*_ parameters and the spiking threshold were the standard ones of the Izhikevich artificial neuron model, whereas the parameters *IZ*_*a*_, *IZ*_*b*_, *IZ*_*c*_ & *IZ*_*d*_ were selected (Table 1, Fig 3E, F) to mimic a regular spiking behavior [31,32]. *IZ*_*input*_ was the input current to the neuron model, that was weighted synaptic activity (*w* ∗*a*) in this article and *IZ*_*k*_ is the input gain factor.

**Table 1.** Izhikevich neuron model parameters used in the evaluation of this study. For the IZ model responses presented in Fig 3E-F and S3 Fig.

| $IZ_A$ | $IZ_B$ | $IZ_C$ | $IZ_a$ | $IZ_b$ | $IZ_c$ | $IZ_d$ | $IZ_k$ |
| --- | --- | --- | --- | --- | --- | --- | --- |
| 0.04 | 5 | 140 | 0.02 | 0.2 | -65 | 8 | 300 |

**Fig 3.**
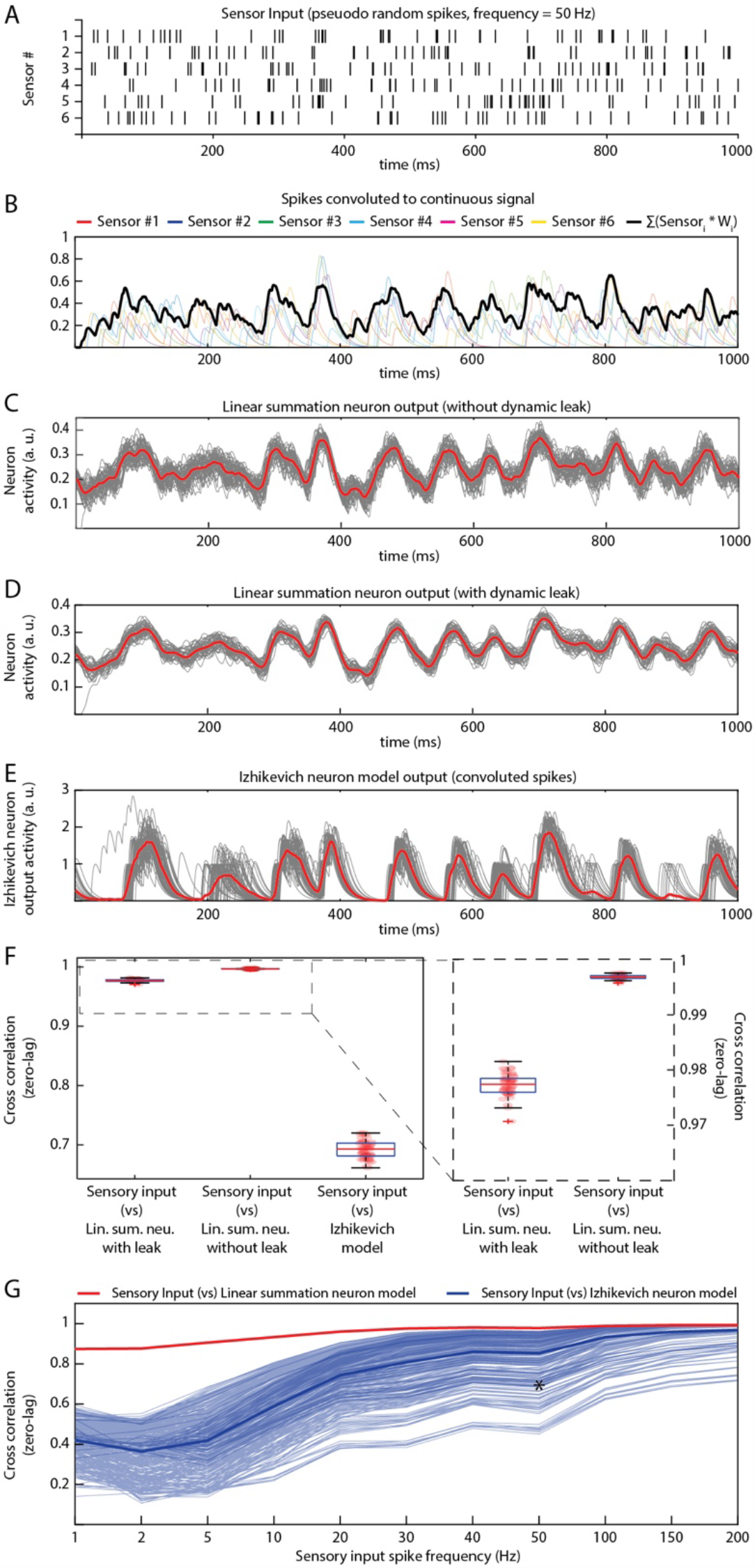
Comparing the properties of different neuron models in isolation. **(A)** Pseudo-random input spike trains (six spike trains corresponding to the six sensory inputs, with an average spike frequency of 50 Hz in each sensor) **(B)** The sensory input spike trains were convoluted using a kernel function (see Methods). The convoluted input responses were fed as weighted (randomly generated, mu = 0.4) EPSP inputs to the neuron model. (**C**) Output responses of the LSM without dynamic leak. The red line is the mean across 50 presentations (each presentation made different by adding Gaussian noise, black lines). In this and all panels below, tests were made for the neuron in isolation, without network connections. (**D**) Similar display as in **C**, but for LSM with dynamic leak. (**E**) Similar display as in **C**, but for output responses of Izhikevich neuron model. The spike output of the Izhikevich neuron model were convoluted using a kernel function (same kernel parameters setting as in **B**). (**F**) Cross-correlations between the sensory input and the output responses of neuron models (illustrated in **C-E**). (**G**) Cross-correlation between different sensory input frequencies and neuron model outputs across a range of IZ model settings, compared to the LSM with dynamic leak. Thin blue lines indicate the cross-correlation with the sensory input for the IZ neuron model responses for each of the 405 IZ model parameter settings (IZ_*a*_, IZ_*b*_, IZ_*c*_, IZ_*d*_ and IZ_*k*_ see methods) tested. Thick blue line indicates the mean of those cross-correlations. The red line indicates the cross-correlation between the sensory inputs and the LSM outputs. Asterisk indicates the cross-correlation measure for the parameters chosen in Fig 3F.

Further, to analyze the IZ model behavior across different spiking and bursting behaviors, we have explored the parametric space (Table 2, Fig 3G) of *IZ*_*a*_, *IZ*_*b*_, *IZ*_*c*_, *IZ*_*d*_ & *IZ*_*k*_ (parameters in Eq. 4-5) within the boundaries identified in Izhikevich, 2003 [31]. We investigated the IZ neuron model responses (Fig 3G) across 405 different parameter settings for each given input spike frequency. The parameter space was defined by the possible combinations of parameters listed in Table 2.

**Table 2.**
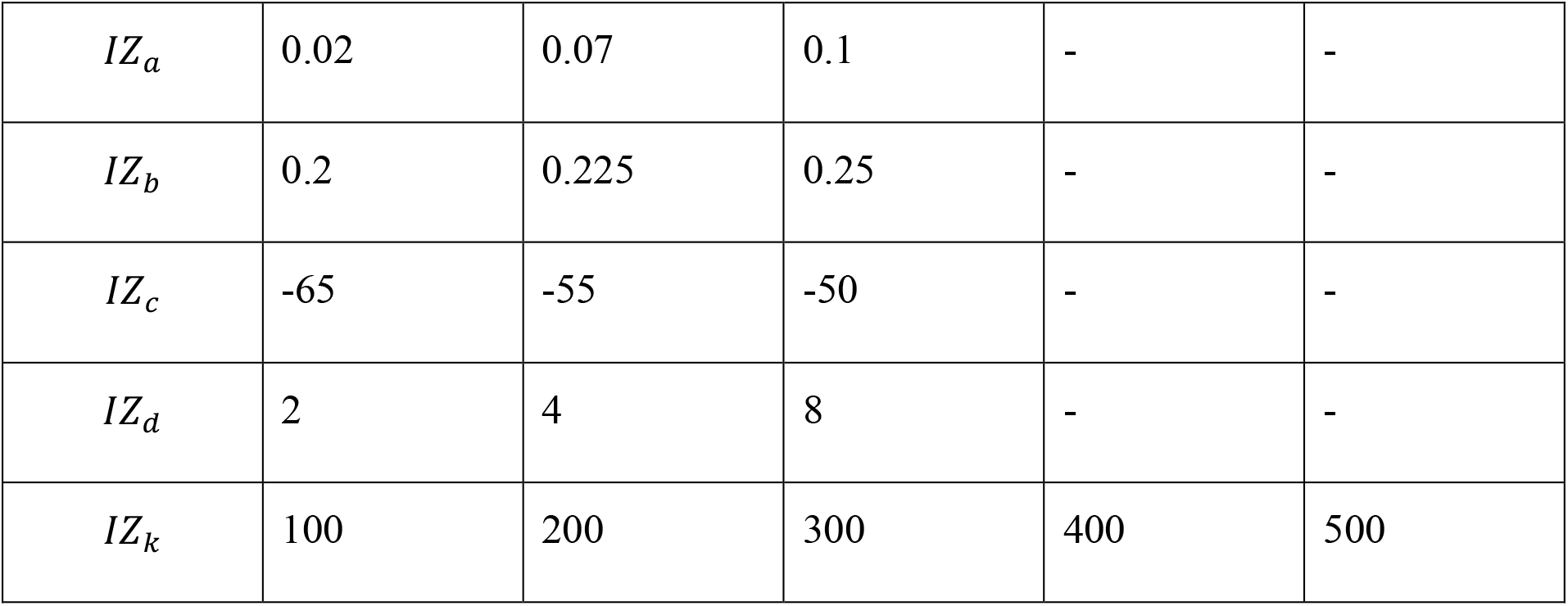
Izhikevich neuron model parametric space explored in the evaluation of this study (for the IZ model presented in Fig 3G).

### Network connectivity

Our network was a two-layer fully connected neuronal network that comprised both inhibitory neurons (*IN*) and excitatory neurons (*EN*) (Fig 4A). The network architecture is defined based on the following two rules: (*a*) The sensory inputs are projected as excitatory synapses to all neurons in layer 1 only; (*b*) All excitatory and inhibitory neurons were fully reciprocally connected both within and between layers. Most of the analysis reported here utilized a “5 × 4” network architecture (5 ENs and 5 INs in layer 1 and 4 ENs and 4 INs in layer 2). In the analysis of Fig 8, where different network sizes were explored, we simply scaled up the number of neurons in each layer using the same connectivity rules (Fig 8).

**Fig 4.**
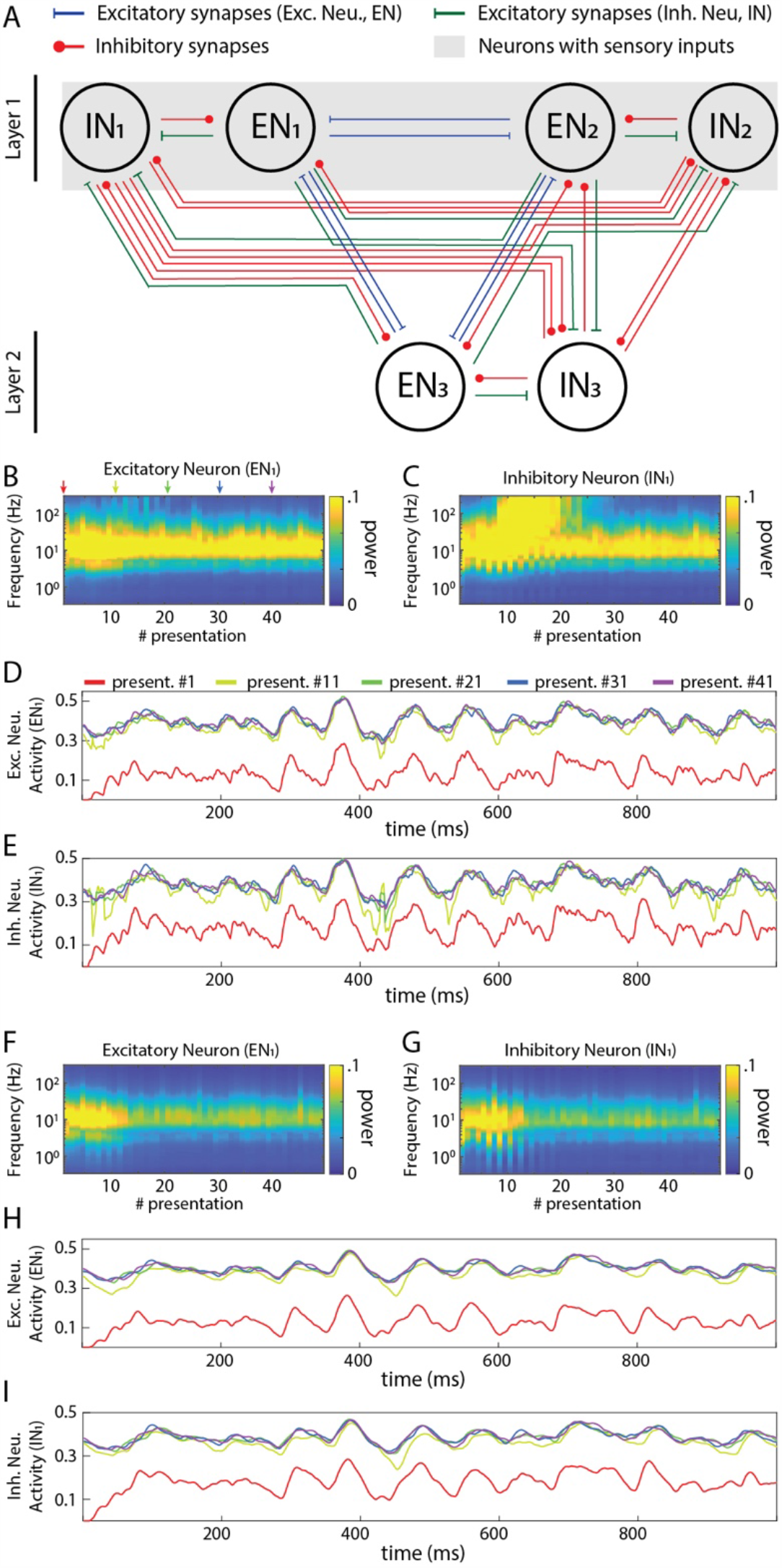
Activity dynamics in a sample recurrent network. (**A**) Principles of the connectivity structure in the recurrent network studied. Note that the default network, and the one used in panels **B-I** of this figure, contained excitatory neurons (5 in the input layer, 4 in the output layer) and inhibitory neurons (same numbers) with the same connectivity whereas only the neurons in layer 1 also received sensory inputs (the same 6 sensory inputs to each neuron) with all synapses having randomly generated gaussian weights (mu = 0.4). (**B**) Frequency plot of the activity in an excitatory neuron. (**C**) Similar plot for an inhibitory neuron. (**D**) Raw data plots for sample signals in the excitatory neuron generated at the indicated presentation #. (**E**) Similar plot for the inhibitory neuron. (**F**)-(**I**) Similar plots as in (**B**)-(**E**) but when all the neurons were modelled to include the neuronal dynamic leak.

A two-layer, fully reciprocally connected neuronal network architecture with self-recurrent connections (autapses) was also investigated. In this specific network architecture, in addition to the network connectivity defined above, the excitatory neurons projected excitatory synaptic connections onto themselves, and inhibitory neurons projected inhibitory synaptic connections onto themselves (S7A Fig).

### Sensory inputs

In this article, we investigated the individual neuron responses (Figs 2, 3 and S1-4 Figs) and network dynamics (Figs 4-8 and S6-7 Figs, except Fig 7C, D) based on six sensory inputs. These sensory inputs were pseudo-randomly generated (see below) and provided as excitatory input to both excitatory and inhibitory neurons. We also tested our recurrent networks with higher input sensor density (#sensors = 6, 15, 30 and 50, Fig 7C, D), the inputs of which were also pseudo-randomly generated.

### Pseudo-random inputs

For the sensory inputs to the LSM and the IZ neurons, we generated pseudorandom spike trains for several different average frequencies (50, 100, 150 & 200 Hz, Fig 3A and S4 Fig) with uniform normal distributions. We used an inbuilt MATLAB® function “*randi”* to generate the spike time distributions in these spike trains. Furthermore, these spikes were convoluted to resemble post-synaptic-potentials (time continuous activity) using the following kernel equation [33],

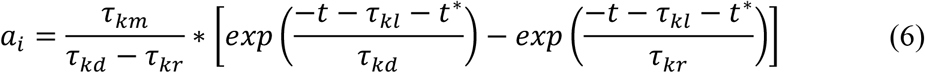

Where, *t*^∗^ is the input spike time, *τ*_*kd*_ is the decay time (4 ms), *τ*_*kr*_ is the rise time (12.5 ms) and *τ*_*km*_ is the constant to calculate ration between rise time and decay time (21.3 ms), and *τ*_*kt*_ is the latency time which is zero in this case. These values were chosen based on the previous work [12]. The convoluted sensor signal was then provided as synaptic input to the neuronal network.

In order to analyze the network dynamics, we provided 50 presentations of the same pseudorandom spike trains (for each given average frequency). Each input presentation differed by an addition of random noise of ± 10 ms to individual spike times (Fig 3C, D, E, black lines) to the reference pseudorandom spike train (Fig 3A, for spike frequency of 50 Hz). These presentations were concatenated without pause or reset between them, so the input subdivided into 50 presentations was in effect one long presentation lasting for 50,000 ms.

To allow a comparison with the output of the LSM, we convoluted the output spike trains also of the spiking neuron model (IZ). The process of convolution emulated a post synaptic response that would have been generated in a receiving neuron, whereas the LSM output itself directly corresponded to such a signal.

### Synaptic weights

All excitatory and inhibitory synaptic weights in the network were randomly distributed, including the excitatory sensory inputs to only the layer 1 neurons. The synaptic weight distributions were either normal, lognormal or binary. The normal and the log-normal distributions were generated for different mean weights (µ) (values between 0.1 − 0.5) each with a fixed coefficiency of variation (*cv*) of 20% (where sigma (*σ*) = (*cv* / 100) ∗ *μ*). For binary distributions, we tested different probabilities of high weight synapses (*w* = 1) (probability varied between 10% − 50%), whereas the remainder of the synapses were set to zero weight (Fig 5A, B, C).

**Fig 5.**
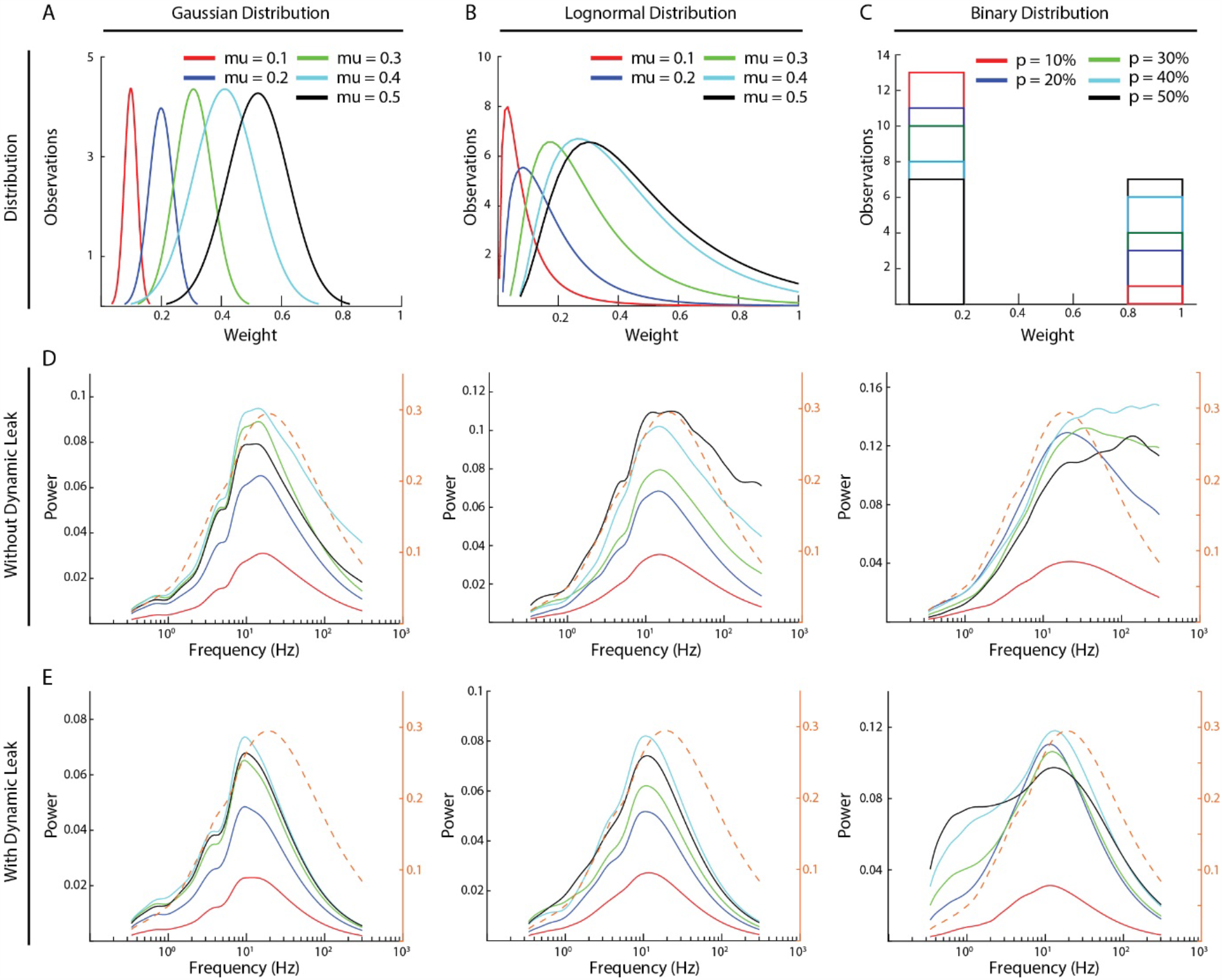
High frequency components and the effect of dynamic leak across different synaptic weight distributions. (**A-C**) The three types of synaptic weight distributions that were explored (Gaussian, Log-Normal and Binary) and the average weight distributions for each mean weight value (five weight distributions were generated for each mean weight). (**D**) The frequency power distributions across all the above synaptic weight distributions. Color keys for the different average synaptic weights are in A-C. (**E**) Similar display as in C, for the same networks but with the neuron model with dynamic leak. Dashed orange traces in D, E show the corresponding frequency power distribution for the sensory inputs at 50 Hz, averaged across the six sensory inputs, for comparison.

### Statistical analysis

#### Cross-correlation

The correlation index measure was used to compute the similarity of the responses of the neuron models (Fig 3F, G and S4E Fig). The correlation between two signals was computed with an inbuilt MATLAB® function “*xcorr”* (with zero lag), which produces values from 0 (uncorrelated) to 1 (identical).

#### Frequency analysis

We performed a continuous wavelet transform (using an inbuilt MATLAB® function *‘cwt’*) in order to define the frequency composition of the input signal over time. The wavelet transform was used to extract the power of each frequency band as a function of time for the continuous neuron activity signal. Here, we reported (Figs 4 – 8 and S5-7 Figs), for each frequency band, the maximum power of the signal within each input presentation time window (1 second).

In S5 Fig, the frequency analysis was performed on the sensory input signals (on the convoluted signal for each given average spiking frequency) across all the 6 sensory inputs for all 50 presentations (see Methods). The maximum power was computed for each sensory input and each presentation, and the average across all 6 input sensors was reported in this figure.

In Fig 4, the frequency analysis was performed on the activity of one excitatory and one inhibitory neuron in layer 1 (Fig 4B, C, F and G) across all frequencies and presentations. S6 Fig display the frequency analysis performed on the activity of all excitatory and inhibitory neurons in the network of Fig 4A. A similar frequency analysis was carried out in Figs 5-8 and S5 Fig, which show the average maximum power calculated across all neurons in all layers and across all 50 presentations.

## Results

### Comparison to Spiking Neuron Models

We first characterized the input-output relationship of the neuron model in isolation (Fig 3) for a standardized sensor input, consisting of randomized spike times in six sensor neurons that were convoluted to time-continuous input signals. These were synaptically integrated by a single modeled neuron (Fig 3A, B). The activity of the non-spiking LSM (linear summation neuron model) was compared with that of a spiking neuron model (Izhikevich, IZ), in terms of how well their output (Fig 3C-E) correlated with the input (Fig 3F). The IZ neuron model was chosen for this comparison, as it was created to mimic a rich neuronal response dynamics with computational efficiency [31]. The spikes generated by the IZ neuron model were convoluted (see Methods) to a time continuous signal (Fig 3E) in order for it to be comparable with the output of the LSM.

Both neuron models (LSM & IZ) were provided with the same pseudo-random sensory inputs (average firing frequency of 50 Hz in each of six sensors, see Methods) connected via six different synapses (Fig 3B). The IZ neuron model parameters for this particular comparison (Fig 3E, F) were chosen to mimic the regular spiking behavior (hypothesized to be a common neuron behavior in cortex [31,32]). The LSM neuron without dynamic leak reproduced on average a close representation of the source convolution signal for the input but the individual traces were considerably noisier without dynamic leak (Fig 3C) than with dynamic leak (Fig 3D). The main difference between the LSM and the IZ neuron model responses was that the IZ neuron model tended to create output dynamic behavior that was not present in the input signal (Fig 3B, E), a consequence of the binary nature of the spike output. A cross-correlation analysis between the neurons’ responses (Fig 3F) showed that the IZ neuron model reflected the input signal less faithfully than the LSM. Note that the cross-correlation is slightly poorer for the LSM with dynamic leak than without, which is due to that the some of the fine-timing details of the high frequency components of the underlying convoluted signal is slightly filtered by the dynamic leak.

We tested if this observation depended on the frequency of the spiking in the sensory inputs. The LSM consistently showed a higher correlation with the input signal than the IZ neuron model across a range of input spike frequencies (S3 Fig). Next we tested if the specific parameters chosen for the IZ neuron model (Fig 3E, also indicated by an asterisk in Fig 3G) were responsible for these results (Fig 3F). Therefore, we tested a range of parameter settings (405 different parameter combinations), which are known to reproduce specific output dynamics (bursting, for example) observed in a variety of neuron types *in vitro* [31]. The correlation analysis showed that LSM was more consistent than the IZ model in maintaining high correlation with the sensory inputs across the full range of sensory input spike frequencies (Fig 3G). The exception was the highest sensory input frequencies, but that can be explained by that the dynamics of the sensory input diminishes due to the density of the inputs (S3A-D Fig), as previously described also for neurons *in vivo* [34]. This effect, which we will refer to as the input density problem, is also evident in Fig 3G.

### Network dynamics (with and without Neuronal Dynamic Leak)

We next investigated the activity dynamics of a standardized recurrent neuronal network implemented using the LSM (Fig 4A). The sensory input was fed as excitatory input to both the excitatory and inhibitory neurons of the first layer for 50 presentations, where the sequential presentation differed by added Gaussian noise to the sensory signal (see Methods). In the network with the neuron model without dynamic leak, there was initially a gradual increase in the power across the higher frequency components of the activity in both excitatory and inhibitory neurons (Fig 4B, C) (more extensively illustrated in S6 Fig, where the first few presentations of sensory input evoked a lower power response). These high frequency components were not present in the sensory input (S5A Fig) and were therefore generated by the network, most likely as a consequence of the parallel excitatory and inhibitory connections, which would be expected to lead to some degree of signal derivation [20]. Interestingly, in the illustrated IN1 the high frequency components gradually built up (until presentation #10, approximately) and then faded away (after presentation #20, approximately), despite that the average intensity of the sensory input did not vary over time, which suggests a relatively rich internal dynamics in this type of recurrent network, despite its limited size. In neurons of the second layer, high frequency components typically faded away more slowly (S6 Fig). The appearance of these high frequency components was sometimes associated with the appearance of transients in the neuron activity (Fig 4D, E). In contrast, in the same network but with the neuron model including the dynamic leak, the transients and the high-frequency components of the neuron activity disappeared (Fig 4F-I). Hence, the low-pass filtering effect of the dynamic leak ‘rescued’ the recurrent network from generating spurious high-frequency components.

The recurrent connections of the network were likely strongly contributing to these high-frequency components. An extreme case of recurrent connectivity is when a neuron makes synapse on itself (autapse). It is not clear to what extent autapses exist in adult neuronal circuitry, but they have been shown to be present in early development [35,36] and they are widely used in the field of RNN/computational neuroscience [37]. To explore the impact of autapses we used the exact same network architecture used in Fig 4 but added autapses to all neurons (S7A Fig). In this scenario, the high-frequency components were strongly amplified (S7B-E Fig). However, in the same network with the neuron model with the dynamic leak, the transients and the high-frequency components of the neuron activity were again effectively removed (S7F-I Fig). We did not explore networks with autapses any further.

We next compared the frequency power distributions of the neuronal activity in this recurrent network across a range of different synaptic weight distributions (Fig 5). We studied three different types of synaptic weight distributions (Gaussian, log-normal and binary distributions). For each type of distribution, we tested five different mean synaptic weights (Fig 5A-C). Moreover, for each given synaptic weight distribution and mean weight, we generated 5 random weight distributions. The average signal of these 5 random weight distributions was used to calculate each frequency power distribution illustrated (Fig 5D, E), where each line represents the average activity across all the neurons of the network (Fig 5D, E). In the network with the neuron model without dynamic leak (Fig 5D), the relative power of the high-frequency components was amplified for synaptic weight distributions at mean synaptic weights of 0.3-0.4 or above (*μ* ≥ 0.4 for Gaussian and *μ* ≥ 0.3 for log-normal distributions) and for p > 10% for binary distribution, compared to the sensory input (S5 Fig). For other synaptic weight distributions (*μ* = 0.1 for Gaussian and log-normal distributions and for p = 10% for binary distributions, for example), there was much lower overall activity in the network, which could be the reason why the high frequency components were not induced in these networks. In the network with the neuron model with the dynamic leak component, the transients and the high-frequency components of the neuron activity disappeared for all settings (Fig 5E), though the setting of the dynamic leak component used also appeared to over-dampen the sensory input dynamics between 20-200 Hz.

To further explore if the high-frequency components observed were induced by the recurrent network, we tested if we could affect the ‘center of gravity’ of the high-frequency components by introducing different conduction delays in signal transmission between the neurons (Fig 6). In the brain *in vivo*, these would correspond to the axonal conduction delays and synaptic delays combined. The delays were randomized between all the neurons, and several different mean delays were tested in different simulations. Interestingly, the peak frequency of the high-frequency component observed without dynamic leak (Fig 6A) was approximately inversely proportional to the mean conduction delay. These peaks were removed by adding dynamic leak (Fig 6B).

**Fig 6.**
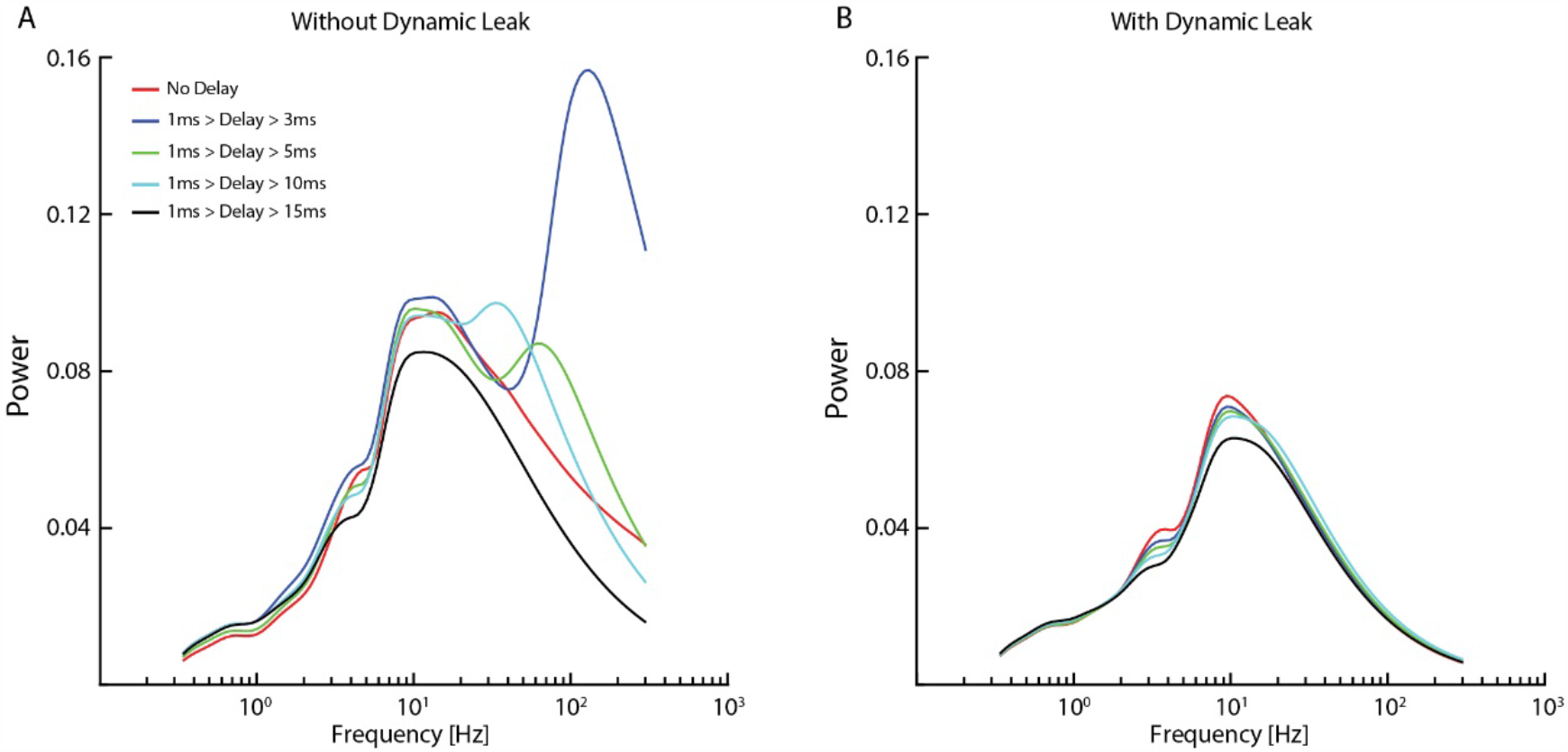
Impact of conduction delays on the frequency distribution. (**A**) Frequency distributions for the same network (network settings as in Fig 4) with different average conduction delays between neurons. (**B**) Data for the same networks and delays when the neuron model included dynamic leak.

**Fig 7.**
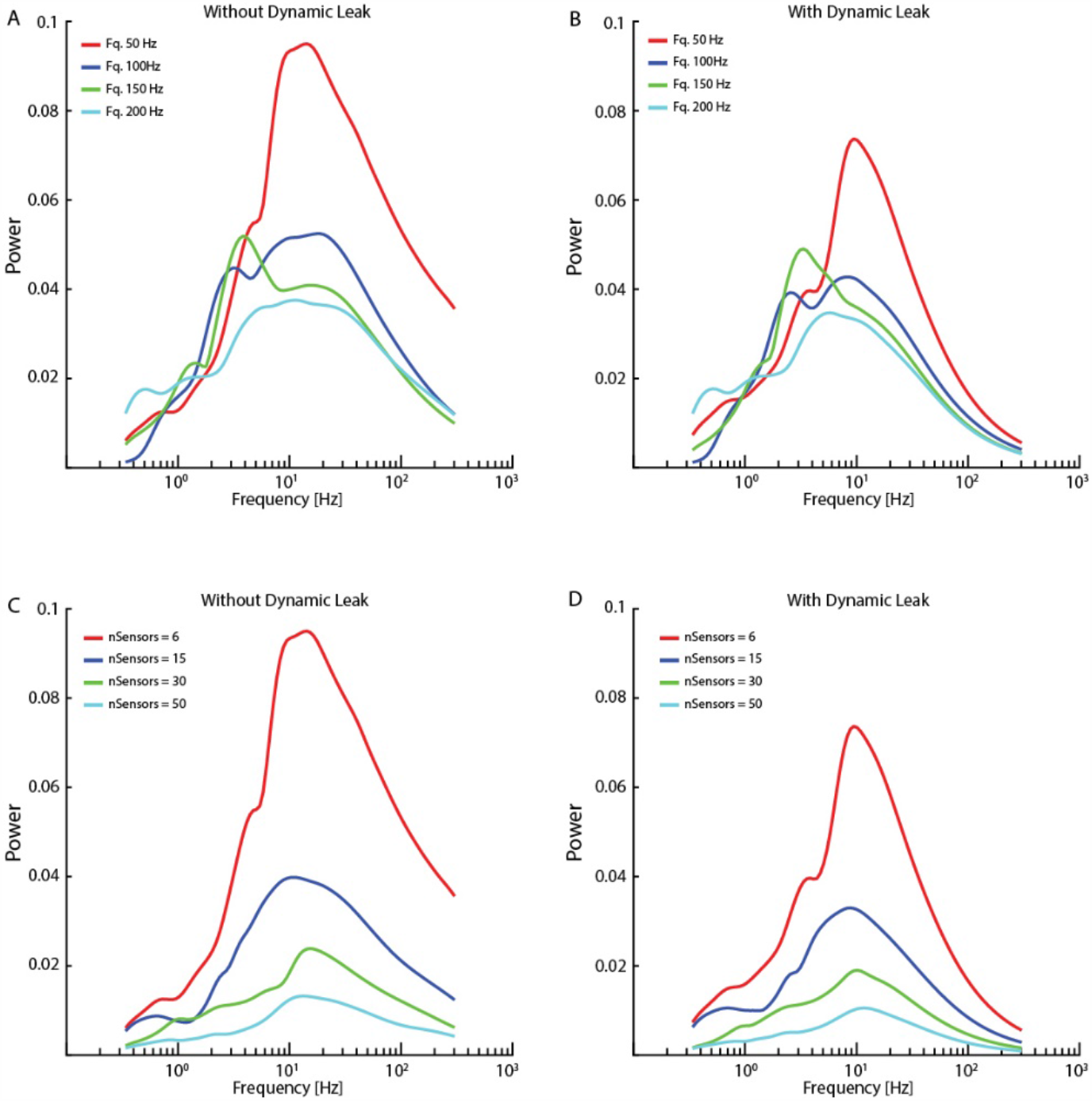
Network activity dynamics increased when sensor input density decreased. (**A**) The high frequency components became more prominent with lower average spike frequencies in the sensor input. (**B**) Neuronal dynamic leak resulted in disappearance of the high-frequency components across all input spike frequencies. (**C**) The high frequency components also became more prominent with lower number of sensory inputs, while a very high number of sensor inputs substantially reduced overall neuron activity dynamics. (**D**) Introduction of neuronal dynamic leak resulted in disappearance of the high-frequency components irrespective of the number of sensory inputs. The network settings for this analysis were similar to Fig 4, except for sensory input.

**Fig 8.**
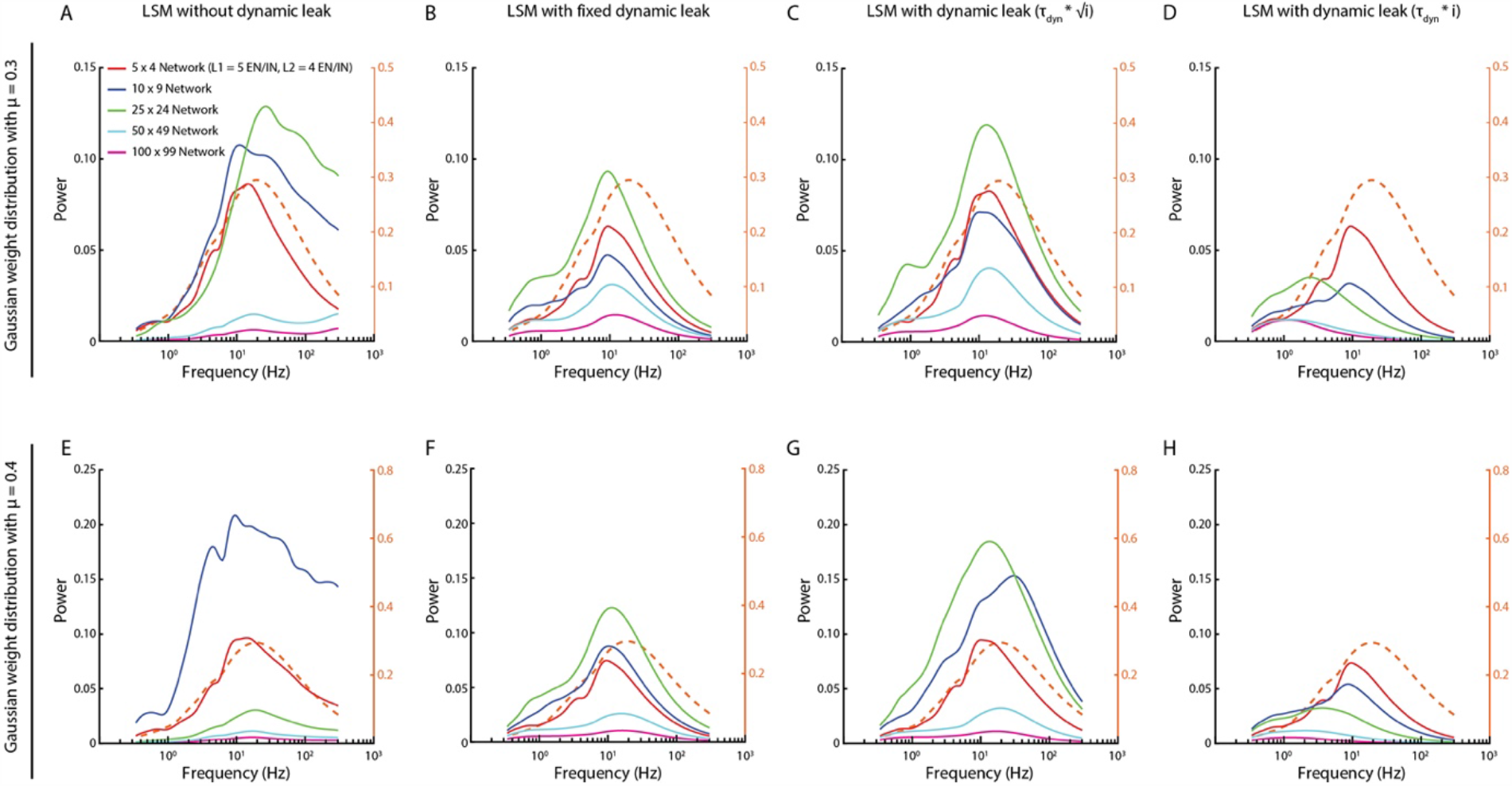
Activity frequency distributions altered with the scale of the networks. (**A, E**) For networks with Gaussian synaptic weight distribution of mean weight mu = 0.3 and 0.4, respectively, the high frequency components could appear without dynamic leak, regardless of network size. (**B, F**) Introduction of the neuronal dynamic leak (τ_JKL_ = 1/100) ‘rescued’ the networks from these high-frequency components. (**C, G**) The dynamic leak constant was adapted based on the square root of number of synapses (*i*). (**D, H**) The dynamic leak constant was adapted based on the total number of synapses (*i*). Dashed orange traces in all plots show the corresponding frequency power distribution for the sensory inputs at 50 Hz, averaged across the six sensory inputs, for comparison.

The high frequency components also appeared for lower average sensor input spike frequency (50 & 100 Hz, Fig 7A) and for lower number of sensory inputs per neuron (nSensors = 6 & 15, Fig 7C). In contrast, for higher input spike frequencies (150 & 200 Hz) and higher number of sensory inputs (nSensors = 30 & 50) the increased density of the inputs resulted in a paradoxical decrease in the power of the neuron activity across all frequencies analyzed (i.e. as shown in S1 Fig), most likely due to the large number of randomized inputs regressing toward the constant mean frequency of each sensory signal. In each case, in the network with the neuron model with the dynamic leak component, the high-frequency components of the neuron activity disappeared for the sensor input configurations where it had been present (Fig 7B, D).

We also explored if the size of the network could be a factor for the appearance of the high frequency components. We found that these high frequency components appeared for different network sizes and that in those cases the network activity was ‘rescued’ when the LSM was implemented with the dynamic leak (Fig 8). Depending on the specific synaptic weight distribution, the high frequency components became unequally dominant for different network sizes (Fig 8A, E) according to unclear relationships. The largest network as a rule had the weakest overall dynamics, which could be due to the same input density problem discussed above, where the density of synaptic input increased as the larger network has a higher number of recurrent synaptic inputs per neuron, which caused the dynamics of the neuron activity to go down. As there is a tendency for membrane time constants to grow with the size of the neuron [30], we scaled the *τ*_*Dyn*_ with the network size (as the neurons of the larger networks had a higher number of synapses) (Fig 8C-D, and Fig 8G-H, for two different weight distributions). A moderate scaling of the *τ*_*Dyn*_ (with the square root of the number of synapses, Fig 8C, G) actually increased the dynamics of some network sizes, while eliminating high frequency components. In contrast, a linear scaling (Fig 8D, H) instead appeared to dampen such dynamics and, unsurprisingly, low pass-filtered also signals well below 100 Hz for the largest networks.

## Discussion

In the present paper, we explored the properties of a non-spiking neuronal model, which was derived from the differential conductance-based H-H model, in various recurrent neuronal networks. We found that in these recurrent networks, many different factors would tend to trigger network induction of high frequency signal components of a somewhat unpredictable magnitude and distribution (i.e. Figs 5-8). These signal components were not present in the input data (S5 Fig) and sometimes peaked to create overt spurious transients (Fig 4B, C). The dynamic leak in our neuron model invariably ‘rescued’ the recurrent networks from their tendency to self-generate these high-frequency signal components (Figs 4-8; S7 Fig). Corresponding to the capacitive component and the ion channels of the membrane circuit, dynamic leak is an inevitable feature of real neurons. Furthermore, this low-pass filter component made the behavior of recurrent networks more predictable for networks of different sizes.

We worked under the scenario that neuronal networks in the brain are recurrent and that excitatory and inhibitory connections are both pervasive, without any a priori assumed structure. Our network architecture contained the circuitry elements of previously reported ‘classical’ network connectivity patterns (feedback and feedforward inhibition, for example). Feed-forward and feedback inhibition running in parallel with excitatory connections was likely the main network feature that caused the signal derivation effects/the high frequency components in the networks without the dynamic leak. The inclusion of autapses in the recurrent network strongly amplified these high frequency components (S7 Fig), presumably primarily through self-amplification of excitatory neurons. But note that in a recurrent network, any local circuity feature will at the global level automatically result in other functional network features as well. Hence, in contrast to a non-recurrent, feed forward neuronal network, in a recurrent network these circuitry features will hence become less clear-cut from a functional point of view, which could cause additional dynamic network effects that for example could explain our observations of gradual build-up of high frequency power components (Fig 4 and S5 Fig) while there was steady sensory input level to keep the network activity up. However, understanding such network dynamics at a deeper level was outside the scope of this paper, but would need to be addressed if such networks are to be used in a functional setting.

In our recurrent networks, apparently spurious high frequency components could be induced for different types of synaptic weight distributions, delays between neurons, sensory input densities and network sizes. It was hard to predict under what exact conditions such high frequency components would become more or less dominant (i.e. Fig 8), but in each case the dynamic leak effectively cancelled them out. From the point of view of the functionality of a processing recurrent network, the fact that the frequency distribution of any given network did not match that of the sensory input is not automatically to be considered a disadvantage because the goal of a processing network would not be to perfectly replicate the sensory input. However, the fact that these high frequency components sometimes took the shape of clear-cut transients with no obvious counterpart in the sensor signal suggests that, at least in part, they should be considered spurious, i.e. noise injected into the signal due to the dynamics of the specific network.

In some cases, the activity of the network became highly suppressed relative to the sensory input (i.e. for low mean weights in Fig 5 and for the largest network in Fig 8). This effect can be ascribed to the input density problem, i.e. when too many unrelated but continuously active synaptic inputs converge on the same neuron, their signal dynamics would tend to cancel out, leaving the neuron with very little signal dynamics [34]. As these signals, due to the network structure, are paralleled by inhibitory connections, when the signal dynamics is lost, inhibition and excitation cancel each other out and the activity dynamics is lost in the network as a whole.

How would spiking neuron networks fare with respect to rescuing a recurrent network from spurious high frequency components? The phasic nature of discrete spike output would be expected to worsen the problem, whereas refractoriness would tend to dampen it. Refractoriness could certainly rescue the system from the extreme transients observed in networks that included autapses. Refractoriness, however, would not rescue the system from high frequency components generated through longer range recurrent excitatory loops.

Recurrent neuronal networks with balanced excitatory and inhibitory synaptic connections have been extensively studied previously [22,23,38,39], using spiking neuron models (employing integrate-and-fire or related mechanisms for the spike generation). In these studies, the recurrent connections were sparsely distributed with an overall connection probability of 1-2%, and a ratio of 4:1 excitatory to inhibitory interneurons. These studies point out that factors such as high connection probability and unbalanced excitation-inhibition tend to produce network instability [39] and in some cases failure in signal propagation across the layers of those neuronal networks [22]. From the stability we observed across a wide range of recurrent network configurations, always at 100% connection probability (though weighted), it would seem that the LSM with dynamic leak would be beneficial for ensuring stable recurrent neuronal network behavior across a range of network sizes and density of connectivity.

The present findings suggest that the biological feature of neuronal dynamic leak, which causes the polarization (i.e. the activity) of the neuron to settle towards resting potential with a time constant, is an important functional feature. It allows brain networks to fully utilize recurrent neuronal network architectures with variable numbers of participating neurons without risking self-generated noise embodied as high frequency components and spurious transients.

## Acknowledgments

Funding: This work was supported by the EU Grant FET 829186 ph-coding (Predictive Haptic COding Devices In Next Generation interfaces), the Swedish Research Council (project grant no. K2014-63X-14780-12-3).

## APPENDIX 1

### Neuron model derivation from H-H model

We first describe the rationale for our Linear Summation neuron model (LSM). In brief, the LSM aims to provide a simple and computationally efficient neuron model, while capturing important characteristics of H-H conductance models [12,40]. The membrane potential in the LSM model is normalized between +1 and −1 with a resting potential of zero. The output of an LSM neuron is a continuous, non-spiking signal that reflects the portion of the membrane potential that exceeds some threshold, which we assumed to be the zero resting potential. This would be suitable to represent one neuron or a population of similarly connected neurons that is biased by background activity to be at or near spontaneous activity (such as is hypothesized for stochastic resonance to prevent dead bands)[41]. This continuous output signal is intended to reflect the mean spike rate that a population of similarly connected neurons would transmit to other centers in the nervous system.

In H-H models the various ion channels associated with ionic pumps and leaks define a resting membrane potential where there are no net currents. Any change in membrane potential away from this resting potential will settle back to the resting potential according to the combined conductance of all of these ion channels, which is called the static leak. Synaptic activation leads to opening of specific ion channels, which in the H-H models as a change of synaptic conductance. The ion(s) that are made permeable by the synapse have a reversal potential that is different from the resting membrane potential. When the ion channels of a particular synapse are open, the membrane potential will be driven towards that reversal potential, with a strength that depends on the strength of the synaptic conductance relative to the static leak conductance.

The synaptic currents charge the neuron, which is modeled as a single capacitor (assuming that the electrotonic distances between different parts of the neuron are negligible). Various synaptic signals are thus integrated and converted into a dynamically changing membrane potential. The static leak is in parallel with this capacitor, thereby defining a time-constant τ for these dynamic changes. The effect is that of low-pass filtering of the integrated synaptic currents [42] to produce the membrane potential that defines the output state of the neuron. In an H-H model, the output state is created by converting the membrane voltage into patterns of spike output, with the help of a threshold for spike generation. Spike generation is omitted in the LSM model and the output of the neuron is instead the part of the membrane potential that exceeds some threshold (herein equal to the resting membrane potential).

First, we describe a conductance-based model similar to previously presented models [12,40] and then show how a model similar to the LSM model can be derived. In this category of ion channel conductance-based models, the dynamics of one type of ion channel is lumped together into one single conductance. Compared to other conductance-based models, no spike generation is modeled and the neurons resting potential is set to zero. The neuron is modeled as a capacitor with capacitance (*C*) and is charged by excitatory (*I*_*syn,exc*_) and inhibitory synaptic currents (*I*_*syn,exc*_) and discharged by a leak current (*I*_*leak*_). Therefore, the neurons membrane potential (*V*) measured across the capacitor follows the following equation.

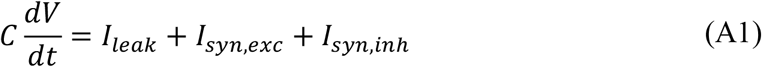

The leak current is set proportional to the membrane potential by constant conductance *g*_*L*_ according to Ohm’s law:

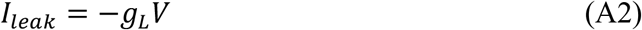

If the synaptic currents are zero, the neuron’s membrane potential will decay to zero. Therefore, this model neuron’s resting membrane potential is zero. At each excitatory synapse *i* the firing rate of the presynaptic neuron 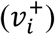 produces a current. The synapse has a baseline conductance 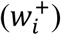, called synaptic weight. This conductance is scaled by the presynaptic neuron’s firing rate. The direction and magnitude of the current is determined by the difference between the constant reversal potential (*E*_*exc*_) and the neuron’s membrane potential. For an excitatory synapse, the current would reverse if the membrane potential rose above the positive reversal voltage, so synaptic activity can never produce a membrane potential about the reversal potential. Therefore if the neuron has already reached this voltage, the neuron’s voltage cannot increase further. The current contributed by an individual synapse can be modeled as

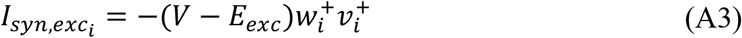

The same model is applied for the current generated by an inhibitory synapse *i*, with reversal potential (*E*_*inh*_), synaptic weight 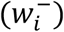 and presynaptic firing rate 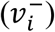. The reversal potential of an inhibitory synapse is set so that the neurons potential cannot decrease below this potential.

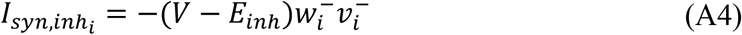

Summing all synaptic currents and plugging in the current equations into the membrane potential equation we obtain

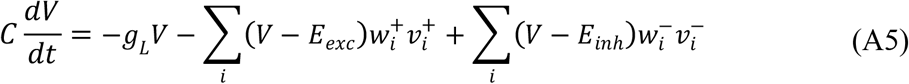

where the first sum is over all excitatory synapses and the second sum over all inhibitory synapses. In order, to show the relationship of this equation to the LSM model, we set the leak conductance to one and replace leak conductance and membrane capacitance by a time constant τ. The reversal potentials are set to +1 and −1 for the excitatory and the inhibitory synapses, respectively. Hence the range of possible membrane potential is between +1 and −1, assuming that the initial voltage is also in that range.

The resulting differential equation is

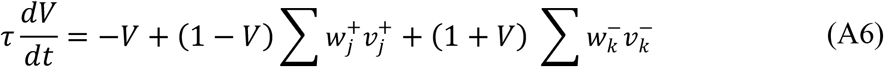

This equation can be solved by applying the implicit Euler method. Let *V*_*t*_ be the membrane voltage in timestep *t* and *h* the stepsize. The following equation must be solved for *V*_*t*+1_

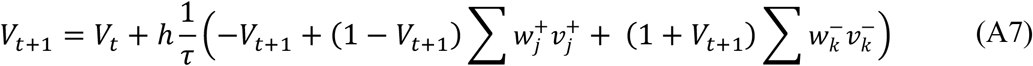

In this case an analytic solution is possible:

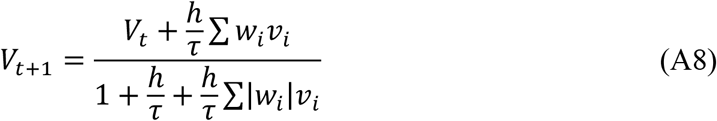

where both sums are over all synapses.

The parameters of this model are also listed in S1 Table. This new system has components not commonly found in neuron models. The reason is that usually the differential equation describes an instantaneous effect of the inputs and the state of the neuron. In this system the effect of the input on how the input is processed by the neuron in a future time step is already considered.

From the above derivation (Eq. A8) we could observe that the excitatory (*I*_*syn,inh*_) and inhibitory (*I*_*syn,exc*_) synaptic currents from a conductance based neuron model (Eq. A1), can be reduced to a total synaptic weight summation 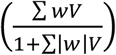, and that the leak current (*I*_*leak*_) (Eq. A1), which is a dynamic leak because it occurs across capacitor, can be reduced to *h*/*τ* (a dynamic leak constant, Eq. A8; if this constant is larger than zero and smaller than one, it can be disregarded in this expression). Based on this neuron model derivation, we propose the simplified linear summation neuron model (LSM, A10) to capture the essential dynamics of the original H-H conductance-based model (S1 Fig). The LSM is given by two equations: LSM without dynamic leak (Eq. A9) and LSM with dynamic leak (Eq. A10).

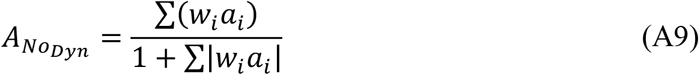

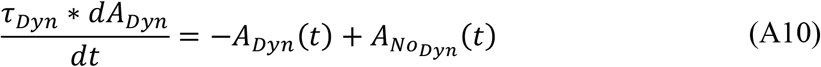

### Supporting Information

**Fig S1.**
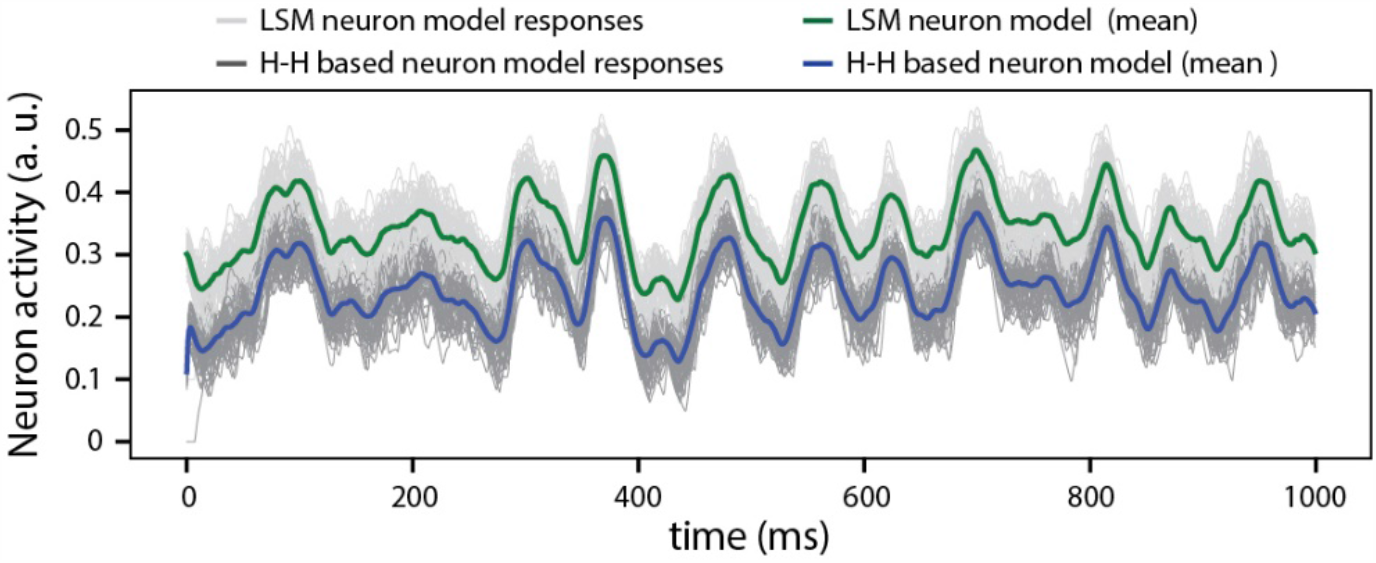
Signal similarity between the LSM and H-H model. A comparison between the output responses for LSM (green line is the mean across 50 presentations) and the H-H (derived using backward Euler method, blue line is the mean across 50 presentations), for a given pseudo-random sensory input at 50 Hz for each of six sensors (see Fig 3). The responses of the LSM output were offset by 0.1 activity (a.u.) in order to visualize the coherence between the responses of both neuron models. The cross correlation (with zero lag) was 0.99.

**Fig S2.**
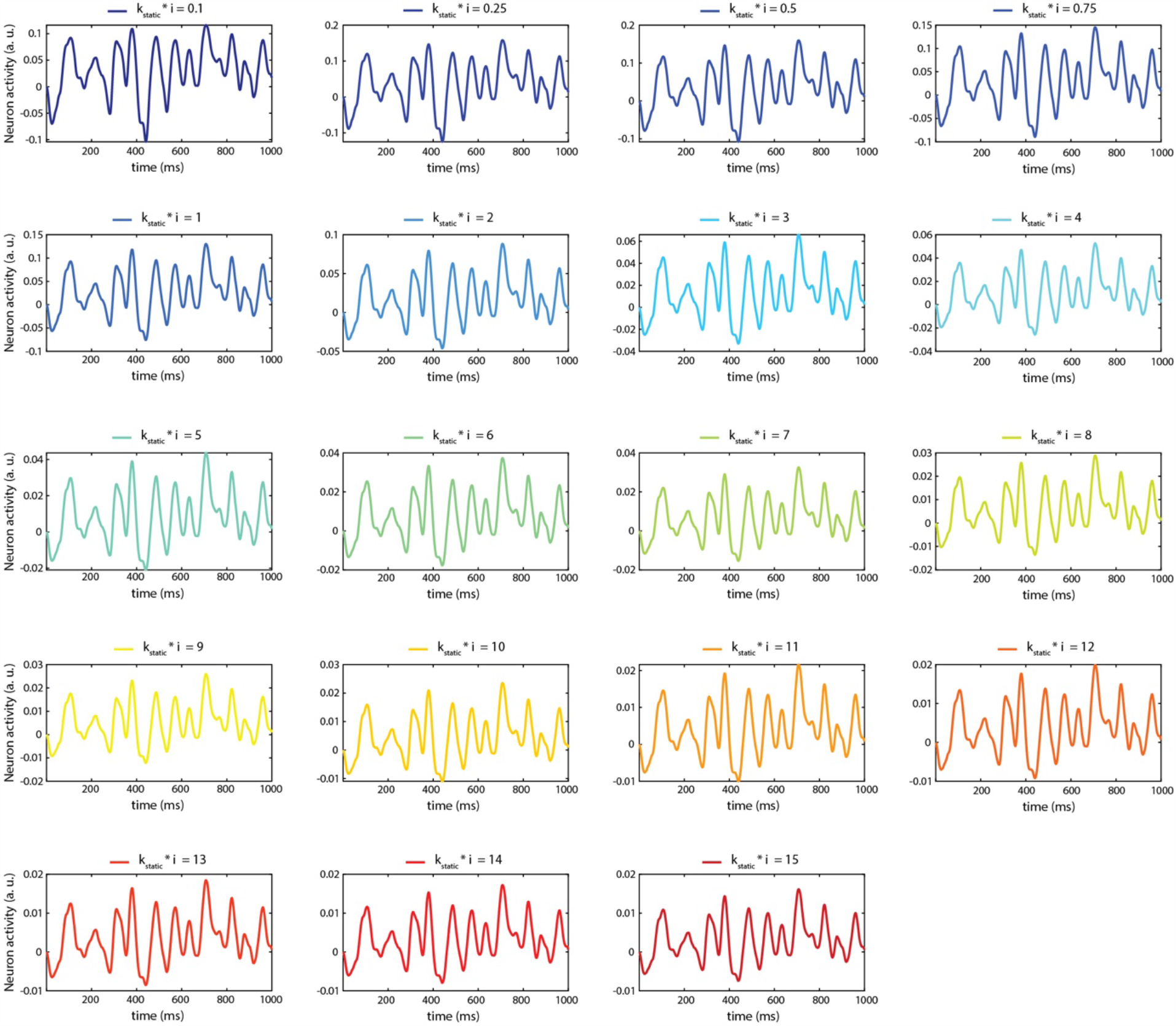
Impact of the value of *k*_*static*_ on the internal activity of the LSM for a given sensory input.

**Fig S3.**
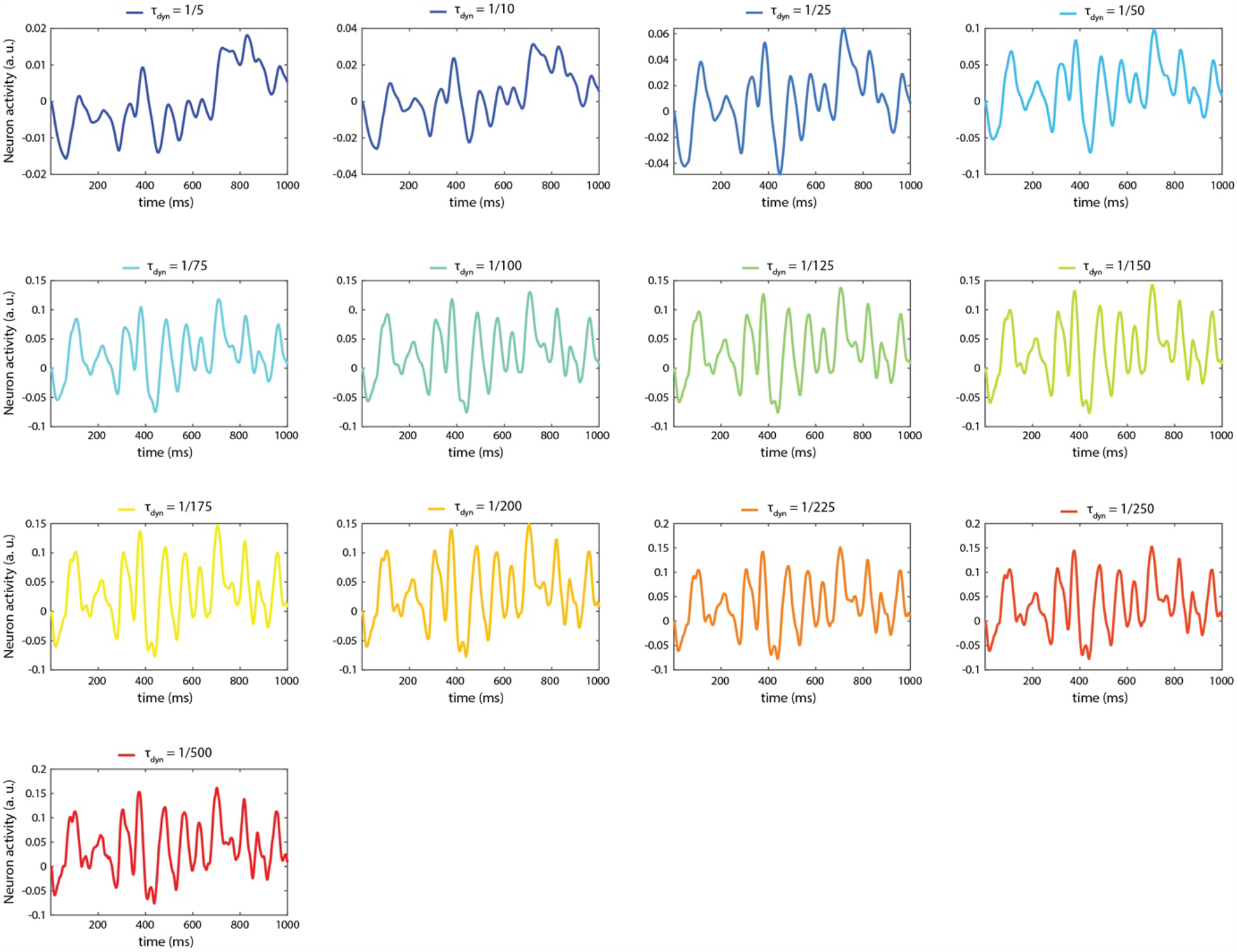
Impact of the value of *τ*_*dyn*_ on the internal activity of the LSM for a given sensory input.

**Fig S4.**
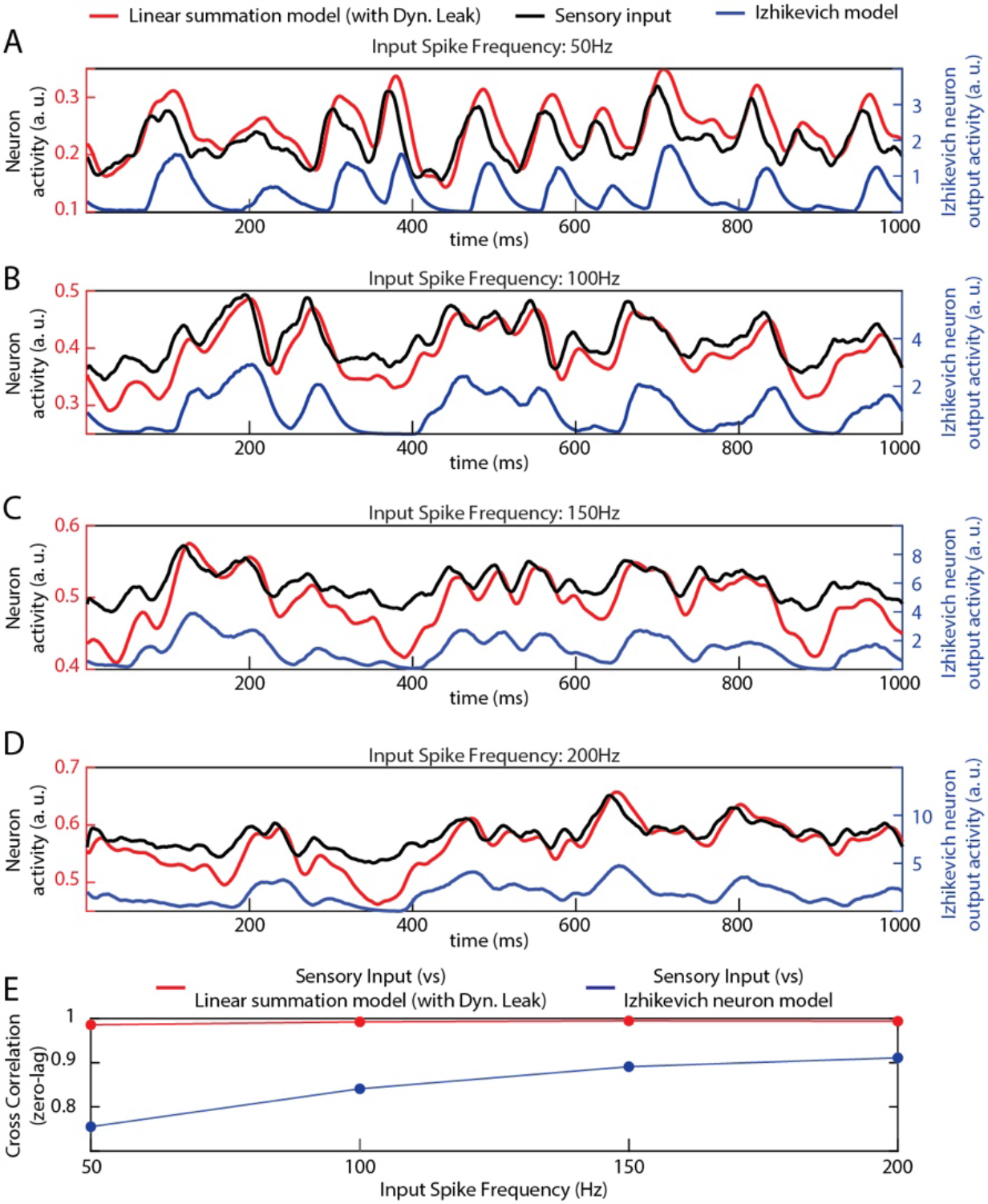
Comparison of the non-spiking and the spiking neuron model outputs for different sensory input frequencies. (**A-D**) Neuron outputs in response to different sensory input frequencies. (**E**) Cross-correlation between sensory inputs and the neuron model outputs.

**Fig S5.**
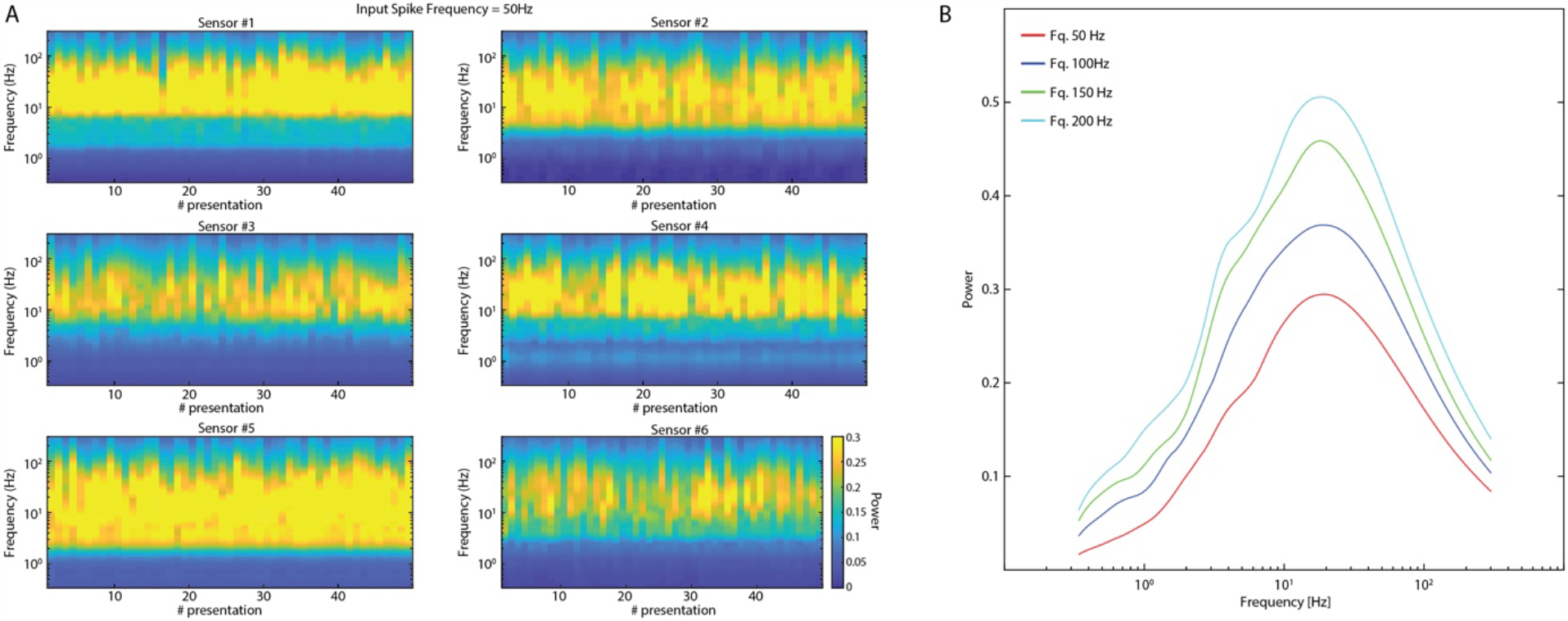
Frequency analysis of the sensory inputs. (**A**) Time-continuous frequency power analysis for each of the six sensory inputs (spike frequency = 50Hz) across the 50 presentations used in the analysis of the network activity. (**B**) Frequency power analysis (using continuous wavelet transform, see Methods), of sensory inputs. The plots show the average power of the activity across all the six sensors, for each of the four mean sensor firing frequencies, across all 50 presentations used in the analysis of the network activity.

**Fig S6.**
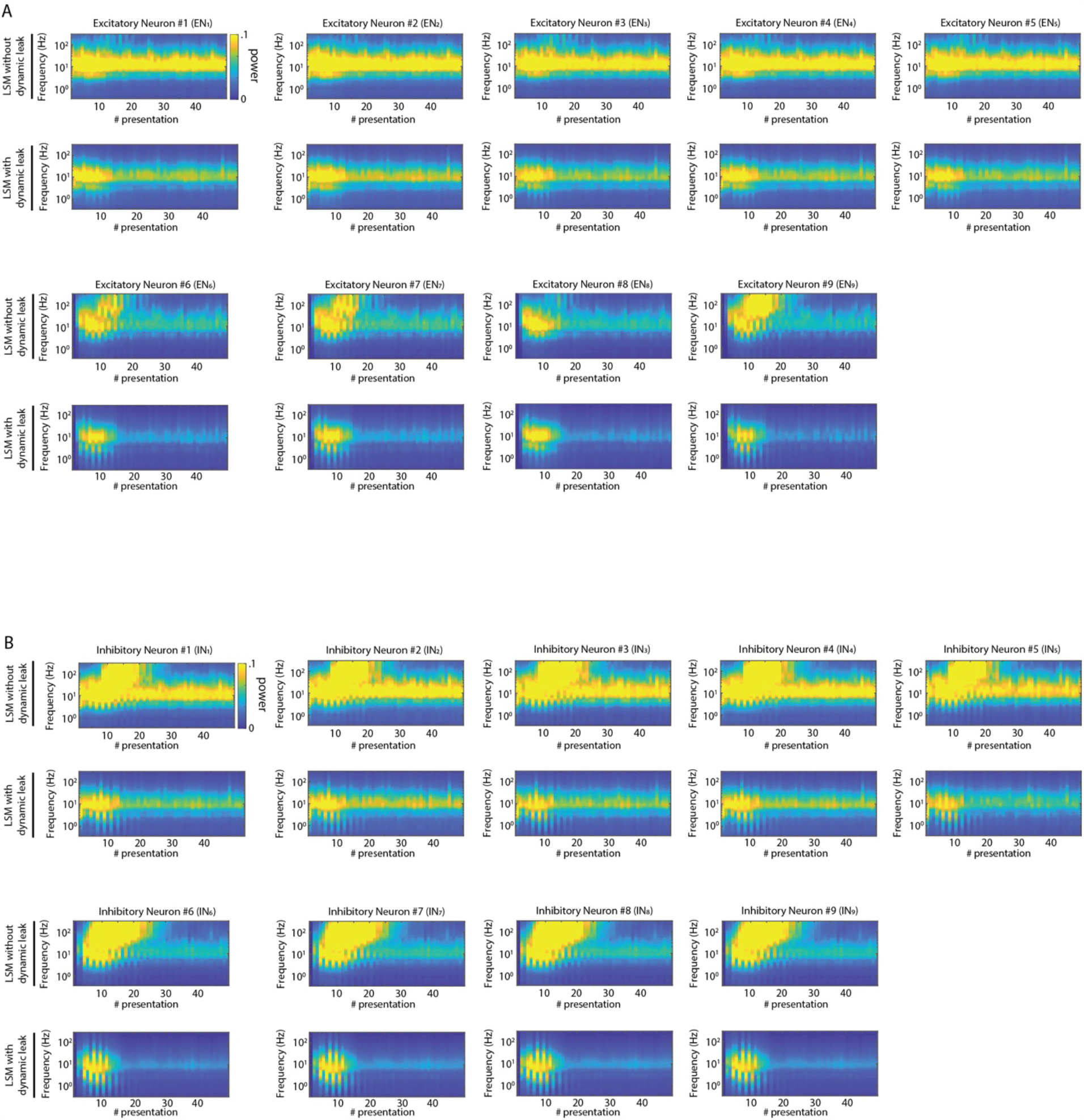
Frequency analysis plots of the activity in all excitatory neurons (*EN*_1_ − *EN*_9_) and inhibitory neurons (*IN*_1_ − *IN*_9_) for the network shown in Fig 4A.

**Fig S7.**
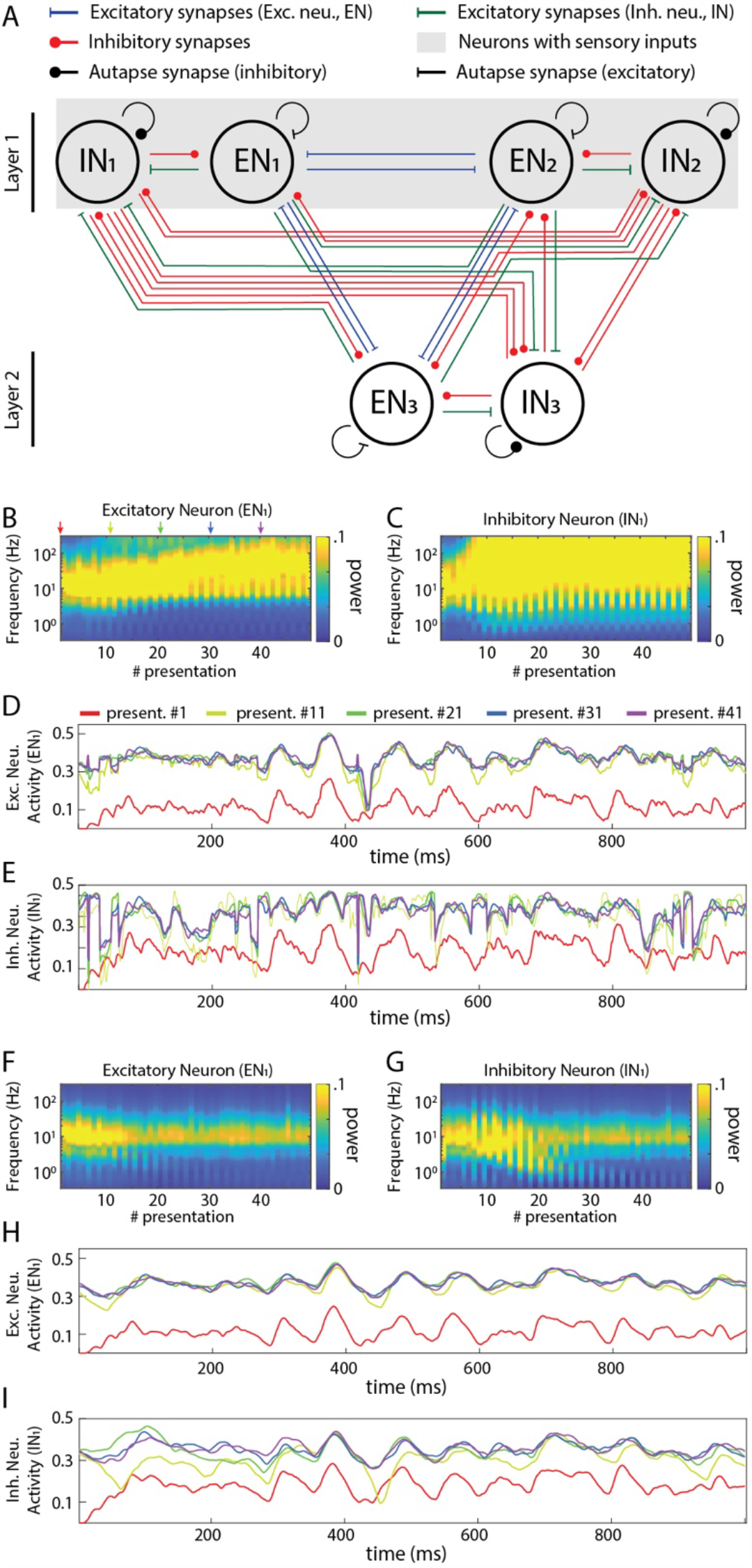
Activity in recurrent networks with autapses. **A**) Principles of the connectivity structure in the recurrent network studied. The network presented here is a fully connected network as in Fig 4, with the addition of self-recurrent excitatory and inhibitory synapses (in excitatory and inhibitory neurons, respectively). (**B**) Frequency plot of the activity in an excitatory neuron. (**C**) Similar plot for an inhibitory neuron. (**D**) Raw data plots for sample signals in the excitatory neuron generated at the indicated presentation #. (**E**) Similar plot for the inhibitory neuron. (**F**)-(**I**) Similar plots as in (**B**)-(**E**) but when all the neurons were modelled with the dynamic leak.

**Table S1.** H-H Model variable definitions (for the H-H neuron model derivation, presented in APPENDIX 1).

| Parameter | Symbol | Units |
| --- | --- | --- |
| Membrane potential | $V$ | [V] |
| Membrane capacitance | $C$ | [F] |
| Membrane time constant | $\tau$ | [s] |
| Time step | $h$ | [s] |
| Neuron output | $v$ | [V] |
| Excitatory weights | $w_i$ | [1] |
| Inhibitory weights | $w_j$ | [1] |
| Presynaptic excitatory neuron activity | $V_i^+$ | [V] |
| Presynaptic inhibitory neuron activity | $V_j^-$ | [V] |
| Excitatory currents | $I_{exc}$ | [A] |
| Inhibitory currents | $I_{inh}$ | [A] |
| Leak currents | $I_{leak}$ | [A] |

## References

1. Binzegger T, Douglas RJ, Martin KAC. A quantitative map of the circuit of cat primary visual cortex. J Neurosci. 2004. doi:10.1523/JNEUROSCI.1400-04.2004

2. Kar K, DiCarlo JJ. Fast recurrent processing via ventral prefrontal cortex is needed by the primate ventral stream for robust core visual object recognition. bioRxiv. 2020. doi:10.1101/2020.05.10.086959

3. Koestinger G, Martin KAC, Rusch ES. Translaminar circuits formed by the pyramidal cells in the superficial layers of cat visual cortex. Brain Struct Funct. 2018. doi:10.1007/s00429-017-1588-7

4. Song S, Sjöström PJ, Reigl M, Nelson S, Chklovskii DB. Highly nonrandom features of synaptic connectivity in local cortical circuits. PLoS Biology. 2005. doi:10.1371/journal.pbio.0030068

5. Hooks BM, Mao T, Gutnisky DA, Yamawaki N, Svoboda K, Shepherd GMG. Organization of cortical and thalamic input to pyramidal neurons in mouse motor cortex. J Neurosci. 2013. doi:10.1523/JNEUROSCI.4338-12.2013

6. Steriade M. Synchronized activities of coupled oscillators in the cerebral cortex and thalamus at different levels of vigilance. Cereb Cortex. 1997. doi:10.1093/cercor/7.6.583

7. Allen GI, Tsukahara N. Cerebrocerebellar communication systems. Physiological Reviews. 1974. doi:10.1152/physrev.1974.54.4.957

8. Jörntell H. Cerebellar physiology: links between microcircuitry properties and sensorimotor functions. Journal of Physiology. 2017. doi:10.1113/JP272769

9. Obermayer J, Heistek TS, Kerkhofs A, Goriounova NA, Kroon T, Baayen JC, et al. Lateral inhibition by Martinotti interneurons is facilitated by cholinergic inputs in human and mouse neocortex. Nat Commun. 2018. doi:10.1038/s41467-018-06628-w

10. Douglas RJ, Martin KAC. Inhibition in cortical circuits. Current Biology. 2009. doi:10.1016/j.cub.2009.03.003

11. Zhu JJ, Lo FS. Recurrent inhibitory circuitry in the deep layers of the rabbit superior colliculus. J Physiol. 2000. doi:10.1111/j.1469-7793.2000.00731.x

12. Rongala UB, Spanne A, Mazzoni A, Bengtsson F, Oddo CM, Jörntell H. Intracellular dynamics in cuneate nucleus neurons support self-stabilizing learning of generalizable tactile representations. Front Cell Neurosci. 2018. doi:10.3389/fncel.2018.00210

13. Swadlow HA. Fast-spike interneurons and feedforward inhibition in awake sensory neocortex. Cerebral Cortex. 2003. doi:10.1093/cercor/13.1.25

14. Isaacson JS, Scanziani M. How inhibition shapes cortical activity. Neuron. 2011. doi:10.1016/j.neuron.2011.09.027

15. Jörntell H, Ekerot CF. Receptive Field Plasticity Profoundly Alters the Cutaneous Parallel Fiber Synaptic Input to Cerebellar Interneurons In Vivo. J Neurosci. 2003. doi:10.1523/jneurosci.23-29-09620.2003

16. Pi HJ, Hangya B, Kvitsiani D, Sanders JI, Huang ZJ, Kepecs A. Cortical interneurons that specialize in disinhibitory control. Nature. 2013. doi:10.1038/nature12676

17. Sultan KT, Shi SH. Generation of diverse cortical inhibitory interneurons. Wiley Interdisciplinary Reviews: Developmental Biology. 2018. doi:10.1002/wdev.306

18. Okun M, Lampl I. Instantaneous correlation of excitation and inhibition during ongoing and sensory-evoked activities. Nat Neurosci. 2008. doi:10.1038/nn.2105

19. Anderson JS, Carandini M, Ferster D. Orientation tuning of input conductance, excitation, and inhibition in cat primary visual cortex. J Neurophysiol. 2000. doi:10.1152/jn.2000.84.2.909

20. Wehr M, Zador AM. Balanced inhibition underlies tuning and sharpens spike timing in auditory cortex. Nature. 2003. doi:10.1038/nature02116

21. Sutskever I, Vinyals O, Le Q V. Sequence to sequence learning with neural networks. Advances in Neural Information Processing Systems. 2014.

22. Vogels TP, Abbott LF. Signal propagation and logic gating in networks of integrate-and-fire neurons. J Neurosci. 2005. doi:10.1523/JNEUROSCI.3508-05.2005

23. Brunel N. Dynamics of sparsely connected networks of excitatory and inhibitory spiking neurons. J Comput Neurosci. 2000. doi:10.1023/A:1008925309027

24. Liou JY, Smith EH, Bateman LM, Bruce SL, McKhann GM, Goodman RR, et al. A model for focal seizure onset, propagation, evolution, and progression. Elife. 2020. doi:10.7554/eLife.50927

25. Chakravarthy N, Tsakalis K, Sabesan S, Iasemidis L. Homeostasis of brain dynamics in epilepsy: A feedback control systems perspective of Seizures. Ann Biomed Eng. 2009. doi:10.1007/s10439-008-9625-6

26. Naundorf B, Wolf F, Volgushev M. Unique features of action potential initiation in cortical neurons. Nature. 2006. doi:10.1038/nature04610

27. Spanne A, Geborek P, Bengtsson F, Jörntell H. Spike generation estimated from stationary spike trains in a variety of neurons In vivo. Front Cell Neurosci. 2014. doi:10.3389/fncel.2014.00199

28. Saarinen A, Linne ML, Yli-Harja O. Stochastic differential equation model for cerebellar granule cell excitability. PLoS Comput Biol. 2008. doi:10.1371/journal.pcbi.1000004

29. Spanne A, Jörntell H. Questioning the role of sparse coding in the brain. Trends in Neurosciences. 2015. doi:10.1016/j.tins.2015.05.005

30. Zhang ZW. Maturation of Layer V Pyramidal Neurons in the Rat Prefrontal Cortex: Intrinsic Properties and Synaptic Function. J Neurophysiol. 2004. doi:10.1152/jn.00855.2003

31. Izhikevich EM. Simple model of spiking neurons. IEEE Trans Neural Networks. 2003;14: 1569–1572. doi:10.1109/TNN.2003.820440

32. Izhikevich EM. Which model to use for cortical spiking neurons? IEEE Trans Neural Networks. 2004;15: 1063–1070. doi:10.1109/TNN.2004.832719

33. Mazzoni A, Panzeri S, Logothetis NK, Brunel N. Encoding of naturalistic stimuli by local field potential spectra in networks of excitatory and inhibitory neurons. PLoS Comput Biol. 2008;4. doi:10.1371/journal.pcbi.1000239

34. Bengtsson F, Ekerot CF, Jörntell H. In vivo analysis of inhibitory synaptic inputs and rebounds in deep cerebellar nuclear neurons. PLoS One. 2011. doi:10.1371/journal.pone.0018822

35. Lübke J, Markram H, Frotscher M, Sakmann B. Frequency and dendritic distribution of autapses established by layer 5 pyramidal neurons in the developing rat neocortex: Comparison with synaptic innervation of adjacent neurons of the same class. J Neurosci. 1996. doi:10.1523/jneurosci.16-10-03209.1996

36. Tamás G, Buhl EH, Somogyi P. Massive autaptic self-innervation of GABAergic neurons in cat visual cortex. J Neurosci. 1997. doi:10.1523/jneurosci.17-16-06352.1997

37. Graves A. Supervised Sequence Labelling. 2012. doi:10.1007/978-3-642-24797-2_2

38. Vogels TP, Sprekeler H, Zenke F, Clopath C, Gerstner W. Inhibitory plasticity balances excitation and inhibition in sensory pathways and memory networks. Science (80-). 2011. doi:10.1126/science.1211095

39. Rubin R, Abbott LF, Sompolinsky H. Balanced excitation and inhibition are required for high-capacity, noise-robust neuronal selectivity. Proc Natl Acad Sci U S A. 2017. doi:10.1073/pnas.1705841114

40. Hodgkin a L, Huxley a F. A quantitative description of membrane current and its applicaiton to conduction and excitation in nerve. J Physiol. 1952. pp. 500–544. doi:10.1016/S0092-8240(05)80004-7

41. Tougaard J. Signal detection theory, detectability and stochastic resonance effects. Biol Cybern. 2002. doi:10.1007/s00422-002-0327-0

42. Lindner B. Low-pass filtering of information in the leaky integrate-and-fire neuron driven by white noise. Understanding Complex Systems. 2014. doi:10.1007/978-3-319-02925-2_22

